# The differential immunological impact of photon vs proton radiation therapy in high grade lymphopenia

**DOI:** 10.1101/2024.06.22.600048

**Authors:** James M. Heather, Daniel W. Kim, Sean M. Sepulveda, Emily E. van Seventer, Madeleine G. Fish, Ryan Corcoran, Nir Hacohen, Theodore S. Hong, Mark Cobbold

## Abstract

Radiation therapy has long been a cornerstone of cancer treatment. More recently, immune checkpoint blockade has also been applied across a variety of cancers, often leading to remarkable response rates. However, photon-based radiotherapy – which accounts for the vast majority – is also known to frequently induce profound lymphopenia, which might limit the efficacy of immune system based combinations. Proton beam therapy is known to produce a less drastic lymphopenia, which raises the possibility of greater synergy with immunotherapy.

In this study we aimed to explore the exact nature of the differential impact of the two radiation modalities upon the immune system. We used multiparametric flow cytometry and deep sequencing of rearranged TCRb loci to investigate a cohort of 20 patients with gastrointestinal tumors who received either therapy. Proton-treated patients remained relatively stable throughout treatment for most metrics considered, whereas those who received photons saw a profound depletion in naïve T cells, increase in effector/memory populations, and loss of TCR diversity. The repertoires of photon-treated patients underwent oligoclonal expansion after their lymphocyte count nadirs, particularly of CD8+ Temra cells, driving this reduction in diversity. Across the entire cohort, this reduction in post-nadir diversity inversely correlated with the overall survival time of those patients who died. This raises the possibility that increased adoption of proton-based or other lymphocyte sparing radiotherapy regimes may lead to better survival in cancer patients.

## Introduction

Radiation therapy (xRT) is a cornerstone of modern cancer treatment; over fifty percent of patients will receive radiation at some point during their care^1^. While it can be extremely efficacious, the primary challenge lies in finding the optimal therapeutic ratio: maximizing the killing of cancer cells while minimizing the dose that normal cells and tissues receive. The last decade has also seen the increased development and application of immunotherapies such as immune checkpoint blockade (ICB), which aim to rally a patient’s own immune cells to recognize and kill their tumors. These agents have revolutionized treatment of several otherwise-recalcitrant cancers, becoming standard-of-care in a variety of cancers and are under investigation in many others, with objective response rates in some tumor types ranging as high as 87%^2,3^.

While radiotherapy and ICB are both successfully treating a broad swathe of patients, their benefits are not seen in all patients or all malignancies. As such many investigators are exploring treating patients with both ICB and radiotherapy, with hundreds of combination trials with tens of thousands of patients undertaken in recent years^4^.

However, it is also well documented that radiation therapy frequently induces lymphopenia in patients undergoing treatment, notably depleting the level of circulating T cells^5^. As these are the very cells that ICB seeks to act upon, the effectiveness of combination RT/ICB trials may be inadvertently blunted. Importantly, in addition to the ramification for combination therapies, radiation-induced lymphopenia is negatively associated with patient outcomes. Severe radiation-induced lymphopenia correlates with poorer prognosis and shorter survival times across multiple cancer types (reviewed in ^6^), independently of histology or prior chemotherapy regimens^7^.

The majority of radiotherapy currently undertaken – more than 99% of patients treated – is photon based^8^. However there is increasing interest in and use of proton therapy, which is known to induce a much more profound lymphopenia than alternative proton-based options^5^. According to the Particle Therapy Co-Operative Group (PTCOG), there are currently at least 136 sites operating proton therapy facilities, with almost 70% of those opened just in the last ten years^9^. Proton therapy allows for more precise delivery of radiation to the target, reducing dose deposition in normal tissue compared to photon-based xRT^8^. While the relative tumoricidal efficacy of proton versus photon therapy is in the process of being determined across a broad range of cancers, a number of studies have already reported a differential effect of the two modalities on lymphocytes. Several studies of patients with esophageal cancer reported a significantly worse lymphopenia produced in photon versus proton therapy, particularly with respect to a greater incidence of severe grade 4 lymphopenia^10–13^.

In this retrospective cohort study, we compared banked blood samples from patients who developed severe lymphopenia following either photon or proton-based radiation therapy, and used high-throughput assays to investigate their immune cell constituents. Given the existing literature, we hypothesized that photon xRT should produce a more profound lymphopenia, corresponding to a less diverse lymphocyte repertoire and a worse recovery of immune cell subsets. We performed multiparametric flow cytometry and T cell receptor (TCR) repertoire sequencing on the peripheral blood of samples before, during, and following lymphoablation to test our hypotheses.

## Methods

### Patients

All deidentified sample donors provided informed written consent, and specimens were collected according to Institutional Review Board-approved protocols in accordance with the Declaration of Helsinki.

All patients detailed in this study were treated between 2016 and 2018 at a single institution, the Massachusetts General Hospital (MGH) Cancer Center. These samples were collected as part of a long running effort to help determine the relative efficacy and considerations of photon-versus proton-based treatments; we have pre-emptively consented patients with various cancer types undergoing radiation therapy and banking samples throughout their treatment. The 20 patients that were the focus of this study were chosen from those who were treated for gastrointestinal cancers, and who had banked blood samples and lymphocyte count data at all three time points under consideration: one at their pre-xRT baseline, another at their lymphocyte count nadir, and a third at a ‘recovery’ time, i.e. post-xRT yet no longer lymphopenic.

The larger cohort of 191 patients used to assess differential lymphopenia were also those with a gastrointestinal cancer (cholangiocarcinoma, pancreas or esophagogastric cancer) who received chemoradiation at MGH, but who were previously radiation naïve. They also required lymphocyte counts throughout treatment, but did not require banked material for inclusion.

Lymphopenia grade definitions were based off absolute lymphocyte counts (ALC) with the following value ranges (expressed in thousand cells per µL):

- 80 <= ALC < 100 = grade 1
- 50 <= ALC < 80 = grade 2
- 20 <= ALC < 50 = grade 3
- ALC < 20 = grade 4

Leukapheresed normal donor peripheral blood mononuclear cells (PBMC) were obtained via from the Massachusetts General Hospital Blood Transfusion Service followed by density gradient centrifugation (Ficoll-Paque PLUS, GE Healthcare) as per the manufacturer’s instructions.

### Treatment

Patients received one of a range of treatment modes based on current best practice, which included intensity-modulated radiation therapy (IMRT), stereotactic body radiation therapy (SBRT), volumetric modulated arc therapy (VMRT), and three-dimensional conformal radiotherapy (3D-CRT). Dose and fraction ranges are specified in Table 1.

### Multiparameter flow cytometry

Blood was collected into Streck tubes (which contain a fixative), before aliquoting and storing at -80°C. While several studies have demonstrated the utility of pre-fixed samples in flow cytometric studies^14–16^, these specific tubes have not been tested to our knowledge. Additionally, peripheral blood leukocytes (PBL) were not separated prior to freezing. We therefore elected to perform flow cytometry using three panels of antibody/fluorophore conjugates, with a variety of partially redundant immunophenotyping markers to allow the maximal recovery of information with embedded sanity checks. Aliquots were thawed and washed twice in FACS buffer (PBS with 2% fetal calf serum and 5 mM EDTA) before being split to three FACS tubes. Cell pellets were then resuspended in one of three panels of antibodies (see Supplementary Tables 1 and 2 for surface markers stained and cell populations derived respectively) and stained for 30 minutes in the dark at 4°C. Cells were finally washed again with FACS buffer and flow data were acquired on a BD LSRFortessa X-20 Cell Analyzer. FCS files produced were analyzed using FlowJo V10.

While multiple stains failed to produce resolvable populations on these fixed cells, many markers were still usable. The frequency of CD4^+^ and CD8^+^ cells was highly correlated across two panels, as were the corresponding CD4:CD8 ratios (Supplementary Figure 6A-C). Similarly, the frequency of CD3^+^ cells in one panel was highly correlated with the sum frequencies of CD4^+^ and CD8^+^ cells in other two panels (Supplementary Figure 6D). Gating on CD4 and CD8 naïve/memory T cell subpopulations (determined by CD45RA and CD27 expression) was confirmed by checking CD57 expression, which should increase across naïve/central memory/effector memory/CD45RA^+^ revertant T cells^17^. Broadly this was observed in our data (Supplementary Figure 6E-F), with the exception of there being a relative decrease in CD57 MFI for CD4^+^ Temra cells, and an increase on CD8^+^ naïve cells.

### T cell receptor repertoire sequencing

One aliquot of frozen blood (harvested from Streck tubes, ∼1.5 ml each) per donor per timepoint was submitted to Adaptive Biotechnologies for gDNA TCRb sequencing on their immunoSEQ platform. These samples were run in October of 2018 using primer set ‘Human-TCRB-PD1x’, to a custom intermediate depth resolution between ‘survey’ and ‘deep’. Primary immunoSEQ data were first converted into an AIRR-seq community compliant standardized format^18^ (making use of proper IMGT-approved TCR gene names) using a custom Python-based tool, `immunoseq2air` (version 1.2.0), available via the DOI <u>10.5281/zenodo.3770611</u> or directly from GitHub (https://github.com/JamieHeather/immunoseq2airr), making use of TCR gene nomenclature from IMGT/GENE-DB^19^. Note that immunoseq2airr was run using the ‘-or’ flag, which suppresses the inclusion of orphon TCR genes (i.e. those situated outside the TCR loci) when there is an ambiguous gene call with at least one non-orphon TCR gene.

### Data analysis

All data were analyzed in Python 3, with the following major shared packages: `scipy` (1.11.4)^20^; `numpy` (1.26.2)^21^; `matplotlib` (3.8.2)^22^; `pandas` (2.1.3)^23^; `seaborn` (0.13.2)^24^. TCR clustering was achieved using `graph_tool` (2.68.)^25^ and `Levenshtein` (0.23.0) packages, while Kaplan-Meier and Cox analyses were performed with `lifelines` (0.28.0)^26^.

### TCR clustering

The top 100 most abundant rearrangements per donor per timepoint were extracted, and their V/J/CDR3 identifiers were pooled and clustered, based off the observation that TCRs which recognize similar epitopes often share sub-sequence motifs and form networks of similar sequences^27–29^. We opted for a stringent clustering method, constructing a graph of TCRs by connecting those which both had matching V and J genes and CDR3 amino acid sequences which matched with an edit (Levenshtein) distance <= 1. In order to ascribe potential antigen reactivities, we used the manually-annotated database of published antigen-specific TCRs, VDJdb^30,31^ (the May 2024 release), filtering out only the human beta chains that: had unambiguous gene calls; began and ended with canonical CDR3 junction ending residues; had a confidence score >= 2. These VDJdb V/J/CDR3s were clustered along with the patient TCRs; any clusters that contained VDJdb-derived sequences with antigens that were >= 90% identical (i.e. same HLA allele, same epitope sequence) were considered markers of potential antigenic specificity for all members of that cluster. Note that data were compared to similar analyses using the antigen prediction tool TCRex^32^, which produced broadly comparable results for the antigens shared by both approaches (data not shown).

## Results

### Photon radiation therapy induced higher grade lymphopenia than proton therapy

In order to determine whether our banked cohort aligned with published descriptions of post-xRT lymphopenia, we compared the lymphopenias of patients undergoing their first course of chemoradiotherapy (chemoRT). While more of these patients received photons than protons, we indeed did see that proton-treated patient absolute lymphocyte count (ALC) nadirs were significantly higher, corresponding to significantly lower grade lymphopenias (Figure 1A and Supplementary Figure 1A respectively).

**Fig 1:**
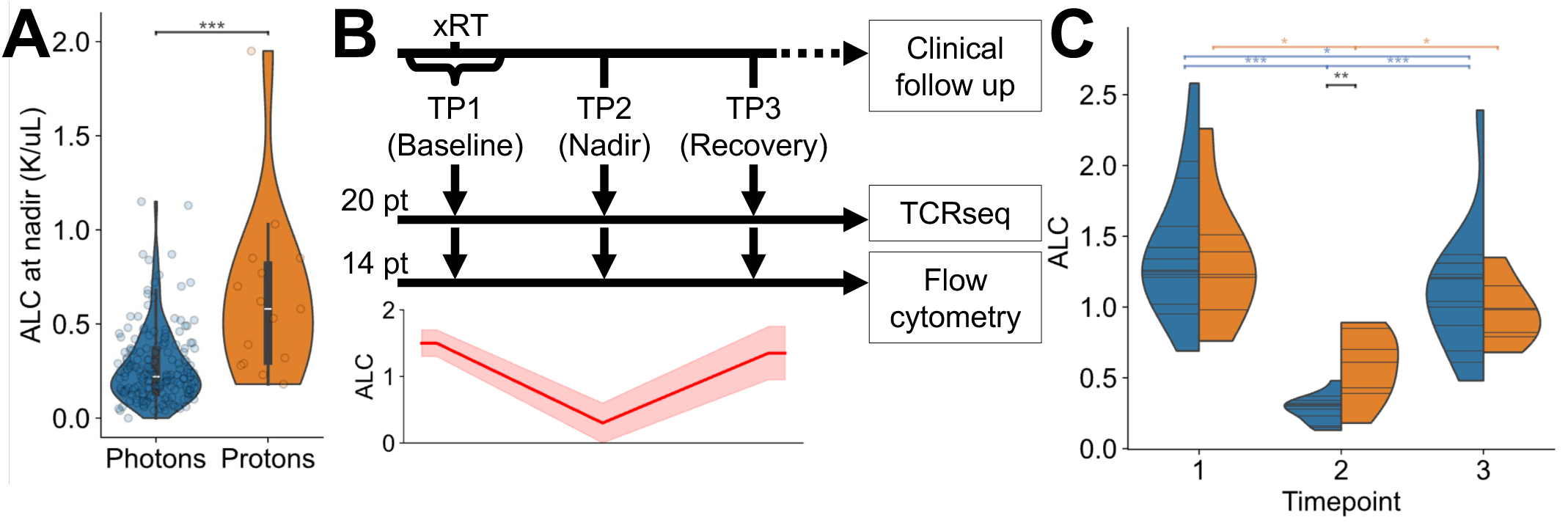
Lymphopenia in the radiation therapy cohort. **A**: Absolute lymphocyte counts at nadir (lowest point following chemoRT) of previously radiation-naïve patients in our wider cohort (n = 190 patients, 175 who received photons and 15 who received protons). ALC expressed throughout in units of thousands of cells per µL. White dots show population medians, thick black bars are interquartile range, and thin black bars are 95% confidence intervals. Violin area shapes indicate kernel density estimations (cut at the terminal observed values). \*\*\**P* < 0.001, Mann Whitney U test. **B**: Schematic of the patient sampling process of the cohort featured in this study. Samples were collected as part of a prospective bio-banking effort. **C**: Violin plots of the absolute lymphocyte counts (ALC) of photon (blue) and proton (orange) treated cancer patients at each of the three time points. Horizontal lines indicate patient values, violin shape indicates kernel density estimations (cut at the terminal observed values). Black significance lines indicate intra-time point unpaired non-parametric tests (Mann Whitney U), while blue and orange lines indicate inter-time point paired non-parametric tests (Wilcoxon ranked-sum). \**P* < 0.05, \*\**P* < 0.01.

These differences were not explained by differences in the nature of the cancers of the patients in each group, as patients in both groups had largely similar cancer types, with the exception of a lack of proton-treated gastric cancer patients (Supplementary Figure 1B).

We then selected twenty patients from the wider cohort who developed high grade lymphopenia over the course of treatment (see Methods) for whom we also had samples prospectively collected. Cohort details are shown in Table 1. These patients required deposited peripheral blood leukocytes (PBL) samples for all three timepoints (TP) of: baseline (TP1), around the time of radiation therapy commencing; nadir (TP2), when ALCs were lowest, and; subsequent recovery (TP3), when ALC values had returned to baseline or otherwise stable levels. PBL for each timepoint from 16 of the donors underwent immunophenotyping via multiparametric flow cytometry, and samples from all 20 donors were processed for T cell receptor (TCR) receptor sequencing (Figure 1B).

Due to the comparative rarity of proton-treated patients with high-grade lymphopenia and the different applications of xRT, the patients selected in this manner were not evenly distributed with respect to cancer and treatment type or duration (Supplementary Figure 2A). While the photon and proton groups were matched with respect to sex ratios (Supplementary Figure 2B), the patients who received proton radiation were significantly older (Supplementary Figure 2C). Despite this difference in age we saw no significant different in ALC between the groups at baseline; however photon-treated patients reached a significantly lower nadir ALC than those treated with protons (Figure 1C), returning to equivalent levels at the recovery timepoint. However, while both groups saw a significant change in ALC from baseline-to-nadir and nadir-to-recovery transitions (more significantly so for photon-treated patients), only photon-treated patients had a significantly lower ALC at recovery relative to their baseline, which did not occur as a result of difference lengths of time between samples (Supplementary Figure 2D). Similarly, it is unlikely that the chemotherapy components of the patients’ treatments influenced our results, as different regimens were adopted approximately equally across both groups (Supplementary Figure 2E). Therefore in this smaller cohort photon-treated patients underwent a larger lymphocyte population contraction and rebound than proton-treated patients.

### Immunophenotypic analysis of lymphocyte population restructuring

In addition to the blood drawn for gathering clinical metrics, additional tubes were taken and banked at each timepoint, where available. 16 of the 20 patients had sufficient banked blood for immunophenotyping by flow cytometry at each of the three timepoints, allowing a more granular analysis of lymphocyte population changes (see Methods for details, Supplementary Tables 1 and 2 for antibody/fluorophore panel information, and Supplementary Figures 3-6 for gating and verification information).

The percentages obtained from the flow data were used to calculate absolute cell numbers using the ALC values described above. This allowed us to observe that T cells were depleted relative to baseline following photon treatment both as a percentage and as a calculated cell number (Figure 2A and B respectively). The reduction in T cell levels from baseline to nadir was not significant in proton-treated patients, although their subsequent recovery was. The nadir reached was also significantly lower for photon-treated patients compared to proton-treated for both measures. We also note that while the frequency of some T cell subpopulations was unchanged (e.g. NKT cells, Figure 2C) the frequency of Treg cells in photon-treated patients at recovery was significantly higher both than it was in the same patients at baseline, and in comparison to the proton-treated patients at the same timepoint (Figure 2D). Other lymphocyte populations were affected, albeit not as dramatically as T cells. For example, B cells were largely stable in frequency across the timepoints in both treatment groups, with a significant reduction to nadir in the photon group and increase in recovery in both (Supplementary Figure 7A and B).

**Fig 2:**
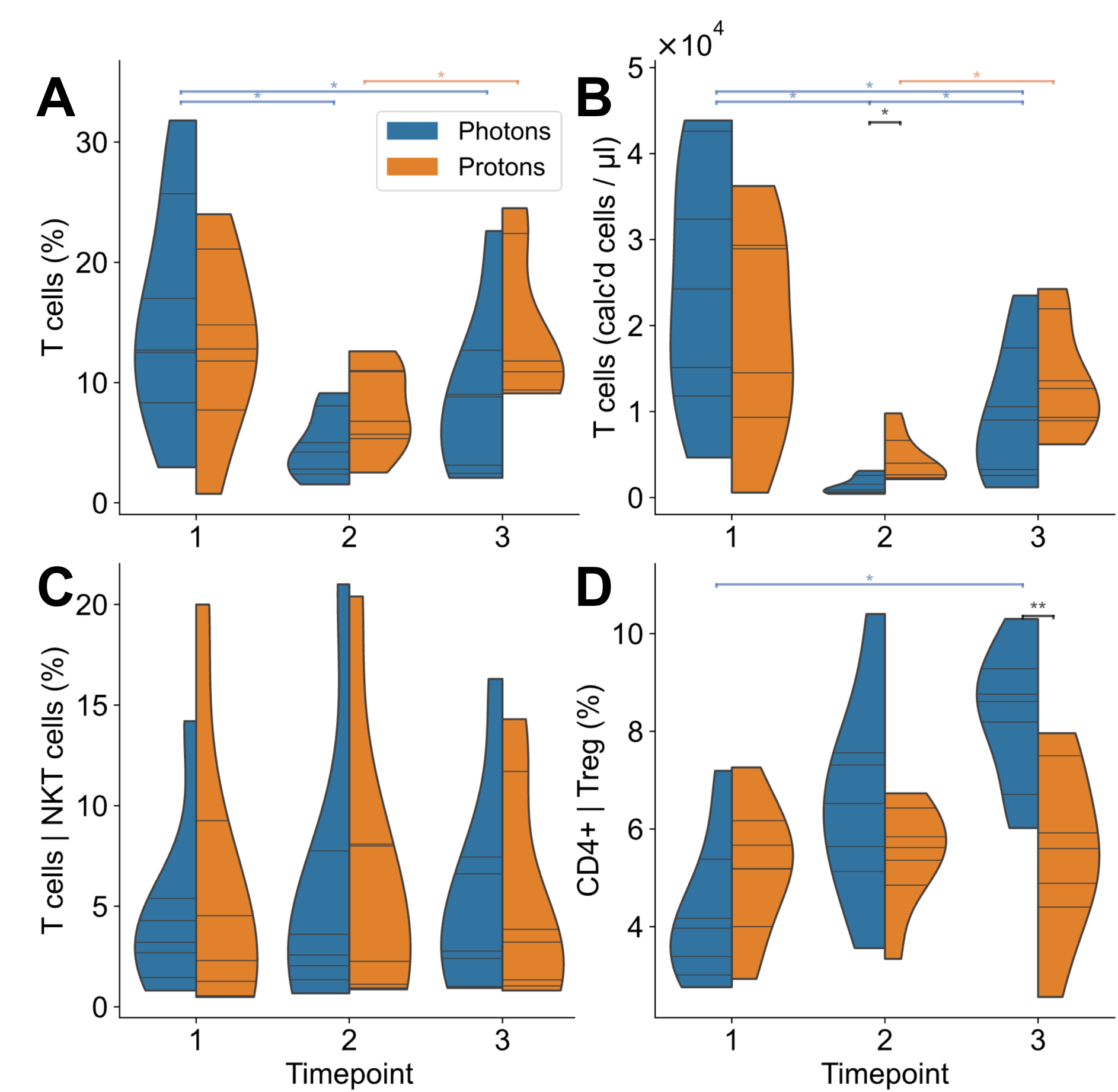
Immunophenotyping of major lymphocyte populations. **A**: Violin plots of the percentage of CD3+ T cells in blood samples of photon (blue) and proton (orange) treated cancer patients at each of the three time points. Horizontal lines indicate patient values, violin shape indicates kernel density estimations (cut at the terminal observed values). Black significance lines indicate intra-time point unpaired non-parametric tests (Mann Whitney U), while blue and orange lines indicate inter-time point paired non-parametric tests (Wilcoxon ranked-sum). \**P* < 0.05, \*\**P* < 0.01. **B**: As in **A**, but showing calculated absolute cell numbers, using the percentage values combined with the corresponding absolute lymphocyte counts (see Figure 1E). **C**: As in **A**, but showing the percentage of NKT cells (CD3+ CD56+). **D**: As in **A**, but showing the percentage of Treg cells (CD4+ CD25+ CD127-).

We then investigated the frequency of T cell subpopulations by differentiation status, looking at CD4 and CD8 naïve (Tn), central memory (Tcm), effector memory (Tem), and effector memory CD45RA+ cells (Temra). Most of the populations remain both stable and equivalent between the two treatment groups across the course of therapy (Supplementary Figure 8). The most notable exception is the CD4^+^ naïve population, which was far more abundant in photon patients at baseline before drastically decreasing on treatment relative to the stable frequencies observed in the proton group, whose low baseline levels are likely explained by being from older donors. CD4^+^ Tem cells displayed a weaker inverse trend (lower in photons at baseline, increasing to equivalent at nadir).

Thus a number of lymphocyte populations are perturbed over the course of radiation therapy, with a greater effect seen in photon patients versus a relatively more stable trend observed in proton therapy.

### T cell receptor sequencing analysis of radiation-induced lymphopenia

In order to assess the potential differential impact of photon versus proton radiotherapy upon the clonal architecture of patient lymphocyte repertoires, equal volumes of blood were processed for beta-chain TCR repertoire sequencing. When taking the abundance (i.e. number of sequencing reads per TCR) into account, we observed that overall photon-treated patients had a significant reduction in TCR-beta rearrangements detected from the baseline to the nadir timepoint, which then significantly rebounded (Figure 3A). The photon nadir samples also had significantly fewer TCR reads than the proton samples, reflecting the pattern observed with ALC values above. When we considered only unique TCRs however (i.e. discounting how frequently each TCR was detected) we saw that there was no significant increase observed among the photon-treated repertoires (Figure 3B), and that both nadir and recovery samples were significantly lower than baseline. In both situations the proton-treated patients showed no significant difference in the number of TCRs between the timepoints, again reflecting the more stable lymphocyte properties observed in the flow cytometry analysis.

**Fig 3:**
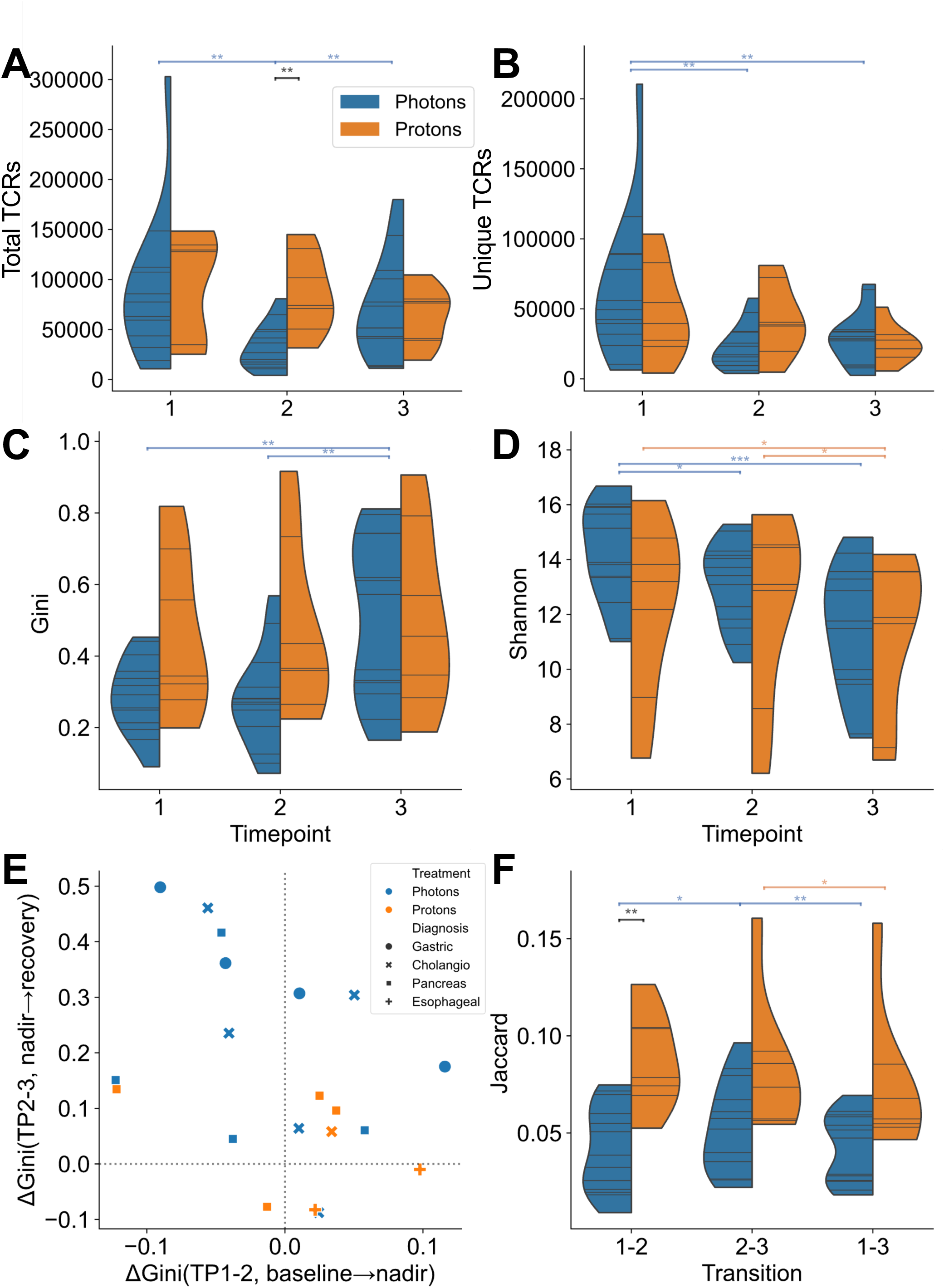
The impact of photon vs proton radiotherapy upon the peripheral TCR repertoire. **A**: Number of total beta chain TCR rearrangements (i.e. factoring in both number of unique TCR rearrangements as well as each of their read abundances) discovered per patient in TCR sequencing of PBL gDNA. Violin area shapes indicate kernel density estimations (cut at the terminal observed values). Black significance lines indicate intra-time point unpaired non-parametric tests (Mann Whitney U), while blue and orange lines indicate inter-time point paired non-parametric tests (Wilcoxon ranked-sum). \**P* < 0.05, \*\**P* < 0.01. **B**: As in A, but showing only the number of unique rearrangements (i.e. ignoring read abundance, counting each sequence only once). **C**: Gini index (effectively unevenness, with values towards 0 being more evenly distributed and those towards 1 being uneven, i.e. oligoclonal) of the beta chain TCR repertoires of the patient samples at each time point. **D**: Shannon entropy (encompassing both species unevenness and richness) of the patient TCR repertoires. **E**: Scatterplot of the change in Gini index of each patient from between timepoints 1 and 2 (x axis) and 2 and 3 (y axis). Samples are colored by treatment type, and markers are assigned by diagnosis. **F**: Jaccard index (a normalized measure of overlap between two sets) of each patient’s whole TCR beta repertoires at each timepoint.

An increase in total TCR read abundance in the absence of a corresponding increase in unique TCR rearrangements suggests that there must be some reduction of diversity of clones present, with some fraction of TCRs in photon-treated patients occupying a greater proportion of the recovery repertoire than at baseline. As such we assessed the patient repertoires using different diversity metrics, which are often employed in such adaptive immune receptor repertoire sequencing (AIRR-seq) analyses^33^, as repertoire diversity is believed to reflect the ability of a repertoire to respond to a wide range of antigens.

The Gini index ranges from zero to one, with zero representing total evenness and one representing total unevenness, which can effectively be treated as a scale of oligoclonality for TCRseq data. Using this measure we saw that while proton-treated patients start off with a more-oligoclonally shifted distribution (higher Gini index) they remain stable throughout (Figure 3C). Photon-treated patients however are only stable between the baseline and nadir samples, with Gini index values at recovery being significantly higher than either previous timepoint. Shannon entropy is another metric that factors in species richness, and thus can be considered a more encompassing diversity metric. Shannon entropies decreased across the timepoints in both treatment groups, albeit with different dynamics (Figure 3D). Photon-treated patients’ baseline samples had significantly higher entropy than the two other timepoints, whereas among proton-treated patients the recovery samples had significantly lower values than the other timepoints. These diversity scores are not an artefact of there being different numbers of TCRs detected per donor per timepoint (largely due to there being different numbers of cells in the equal volumes of blood processed) as randomly subsampling each repertoire to different fixed and equal numbers reveals similar trends (Supplementary Figure 9).

We also visualized the relative stability of proton-treated patient repertoire parameters relative to those who received photons by plotting the change in diversity metrics between the timepoints – ΔGini and ΔShannon – from baseline to nadir (TP1 to TP2), and nadir to recovery (TP2 to TP3), against one another. Figure 3E and Supplementary Figure 10A show that for each metric the proton-treated patients are comparatively localized around zero on both axes, while the photon-treated patients occupy more distant coordinates, highlighting a greater degree of TCR repertoire remodeling across these time periods, especially in their Gini index scores (Supplementary Figure 10B and C). Clinical follow up reveals that the patients’ ALC values are relatively stable post-recovery (Supplementary Figure 10D).

In order to gauge the retention of T cell rearrangements across the course of treatment the Jaccard index (a normalized measure of sharing between two sets) between the three timepoints within each donor was calculated. Figure 3F shows that for whole unsampled repertoires, proton-treated patients share significantly more TCRs between any two timepoints than do photon treated patients. The overlap seen between the nadir and recovery samples (i.e. the transition between timepoints 2 and 3) is significantly greater than between any two other timepoints in photon-treated patients; that is, a TCR observed in the nadir sample is more likely to be observed again in the recovery sample. These properties were again not a product of unequal repertoire depth as they are observed after size-matching via random sampling (Supplementary Figure 11). The TCR repertoires of patients who received photon-based radiotherapy are therefore undergoing more pronounced remodeling events than those who received protons, both at structural and clonotypic levels.

### Correlation of flow cytometric, repertoire, and survival data

In order to see if we could understand the lymphocyte population dynamics underlying the diversity metrics, we leveraged the matched flow cytometry data for those 16 patients who contributed samples to both. To sanity check the principle, we combined samples from both treatment arms and examined their baseline characteristics, which we would expect to most resemble ‘unaltered’ repertoires. We observed that while increased CD4+ T cell frequencies did not correlate with repertoire evenness, increased CD8+ frequencies did correlate with reduced evenness/increased oligoclonality (higher Gini, Supplementary Figure 12A-B). Similarly increased abundance of naïve populations corresponded to more evenness, while increased effector/memory populations corresponded with less (Supplementary Figure 12C-F). This matched our expectations, given that naïve populations are known to be more diverse (more evenly distributed), while CD8+ populations are typically less evenly distributed due to large expansions^34–36^. There was no relationship between the overall CD4+ or CD8+ T cell frequencies and ALC (Supplementary Figure 12G-H).

Some of these correlations are so strong as to be borne out at lower power, after splitting the samples into their treatment groups. Among the strongest correlations are those of the CD8+ naïve and terminally-differentiated Temra populations (Supplementary Figure 13). We observed that at the baseline and nadir timepoints the photon-treated group samples are shifted towards more naïve cells and more diversity relative to the proton group, likely reflective of their initial immune differences (e.g. being younger). By the recovery timepoint however, the photon-treated patient samples have changed to mirror the correlations of those treated with photons, who themselves remained consistent throughout. This property was mirrored in the CD4+ compartment, when looking at naïve and Tem cells, albeit with greater variance (Supplementary Figure 14). We also asked whether the changes in flow and repertoire metrics might better highlight responsible parties. By far the greatest correlation observed was between the change in photon-treated patient CD8+ Temra cell frequencies and whole repertoire TCR diversity in the nadir-to-recovery transition (Figure 4A and Supplementary Figure 15). It therefore appears that the dramatic remodeling of the T cell compartment in photon-treated patients who developed lymphopenia is driven by terminally-differentiated CD8+ T cell expansion post-nadir. This remodeling was also visualized by plotting the frequencies of the largest rearrangements across the timepoints, revealing individual TCRs taking up a far greater proportion of the recovery repertoires (Figure 4B and Supplementary Figure 16).

**Fig 4:**
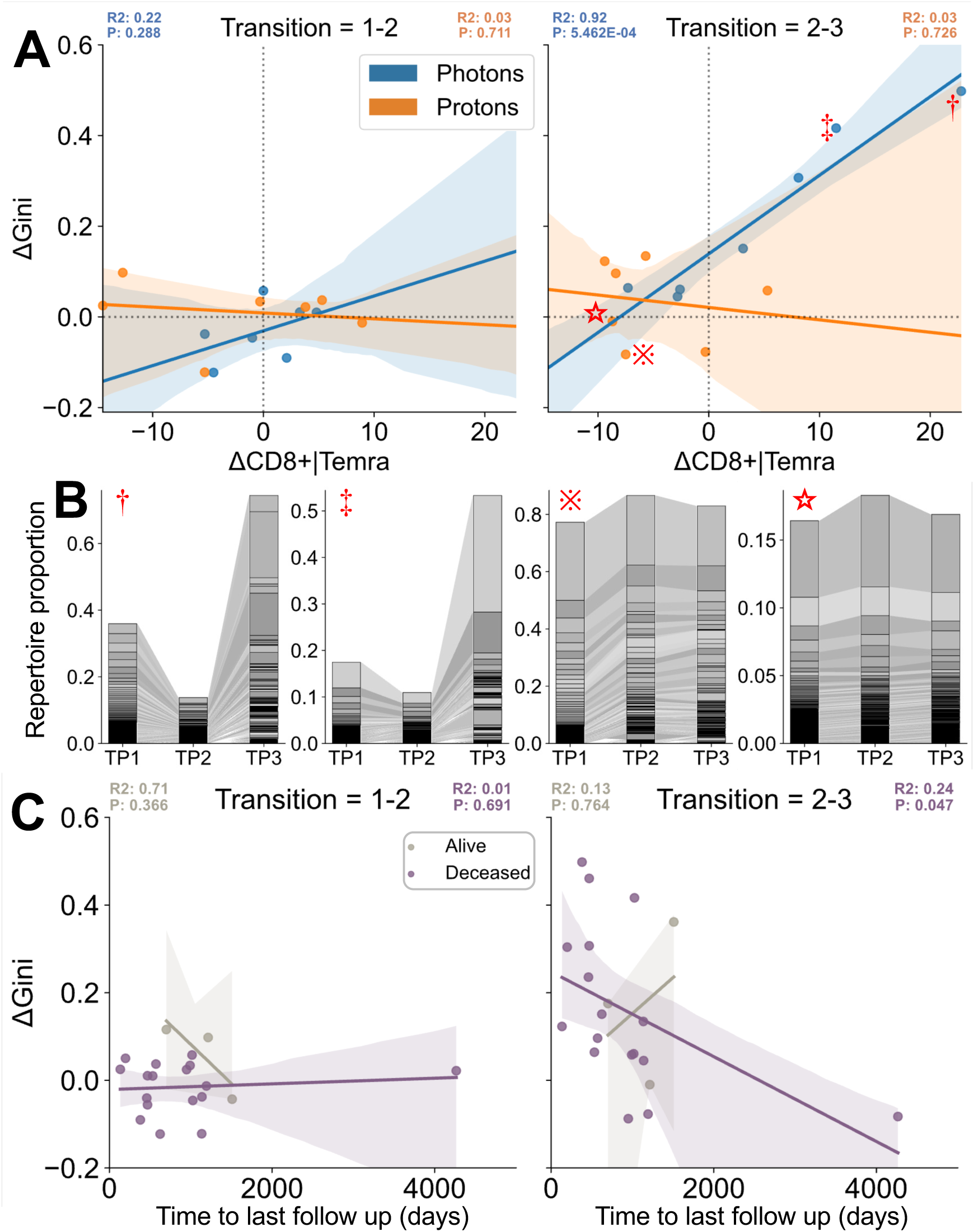
Correlation of cytometric vs repertoire T cell parameters. **A**: Linear regression of the photon (blue) and proton (orange) treated patient samples, showing the inter-timepoint change in percentage frequency CD8+ Temra cells (CD8+ CD27-CD45RA+) on the x axis and change in Gini index of size-matched TCR repertoires (average of sampling 4000 TCRs 100 times) on the y, for each time point. Shaded areas indicate 95% confidence intervals. Color-matched R^2^ and P values displayed above show each patient group’s regression statistics. **B**: Example Sankey-style flow plots showing the change in frequency of the most abundant TCR rearrangements in select donors, keyed to their position on the right-hand panel of **A**. The top 100 rearrangements per donor per time point were pooled, assigned random greyscale colors, and plotted in stacked bar-charts with connecting shaded areas (with absence in a timepoint indicated by shaded areas originating from the halfway point between stacks). Left-to-right the donors involved are: TPS210 (†); TPS109 (‡); TPS219 (**※**); TPS204 (**⋆**). **C**: Linear regression of the patients who were alive as of the last sampling (grey) and who died (purple) in the course of this study, showing change in Gini index on the x axis versus time from diagnosis on the y. Left plot shows the transition from baseline to nadir, middle plot shows nadir to recovery, right plot shows baseline to recovery (skipping nadir). Shaded areas indicate 95% confidence intervals. Color-matched R^2^ and P values displayed above each plot show the relevant patient group’s regression statistics.

We performed clustering of the top 100 most frequent clones from each time point of each donor, along with TCRs of known specificity manually annotated from the literature (via the VDJdb resource)^30,31^, to see if we could identify potential candidate antigens driving these large expansions. While at the cohort level there appeared to be an increase in potentially cytomegalovirus (CMV)-reactive rearrangements in the nadir-to-recovery transition (Supplementary Figure 17A), closer inspection revealed that most donors had little evidence of consistent matching to specific antigens. There were two donors which had TCRs clustering with multiple epitopes restricted by the same HLA allele, each displaying a post-nadir increase in putative CMV-reactivity (Supplementary Figure 17B-C). However of these two, one lacked accompanying flow cytometry data and the other was not a patient that underwent a post-nadir loss of diversity or CD8+ Temra expansion, so they do not illuminate which antigens might be driving the large expansions in the photon-treated cohort.

In order to assess whether these repertoire dynamics have any relationship with patient outcome we plotted the change in Gini index between the different timepoints against time from diagnosis until last follow up. Of the 20 patients, 17 died across the data collection timeframe. While there is no relation for the ΔGini from baseline to nadir (Figure 4C, left), the change in Gini index between nadir and recovery samples (right) marginally but significantly correlates with survival time (P = 0.047). Thus a large decrease in repertoire evenness – or increase in repertoire oligoclonality – following xRT treatment associated with shorter survival in this cohort. However Kaplan-Meier and Cox model analyses of the post-nadir Gini index changes did not find any significant difference (Supplementary Figure 18) between patients with marked increase or decrease in diversity post-nadir, despite a separation of the curves, indicating that we are underpowered to draw strong conclusions of how post-nadir T cell repertoire remodeling influences patient survival.

## Discussion

In different clinical settings, radiation and immune checkpoint blockade therapies individually have been shown to be effective tools in the anti-cancer arsenal, driving huge interest in finding optimal combinatorial strategies. However as radiation therapy can ablate large numbers of the very immune cells required to be activated for successful immune checkpoint blockade, there is a need to better understand its impact upon the immune system, so that those combinations can be rationally designed. As such we have undertaken a comparative study of human T cell population dynamics following either photon-or proton-based radiation therapy. Longitudinal blood samples from prior to radiation, from the absolute lymphocyte count (ALC) nadir, and from a subsequent date when the ALC had recovered, were drawn from 20 patients who received either form of treatment. Samples from 14 of those patients were processed with multiparametric flow cytometry, and samples from all 20 patients had their bulk TCR beta chain repertoires sequenced, revealing comparatively more dramatic T cell remodelling in the photon-treated patients.

One limitation of this study is that the input treatment cohorts were not perfectly matched with respect to cancer types and age. This likely reflects the fact that different tumor types occur across different age ranges, and current clinical practice is likely to direct certain tumors to one treatment modality over another. As such we observed that the photon-treated patients tended towards more diverse and less differentiated T cell repertoires at baseline, likely largely due to them being younger on average than those who received protons. However, despite this initial difference we observed that by most metrics considered photon-treated T cell compartments underwent the most dramatic changes. Their ALC contracted and rebounded more drastically; their T cell frequencies decreased more and did not recover to the same extent; they saw a large shift from naïve to effector memory phenotypes, and their repertoires became far less diverse and more unevenly distributed. This pattern is highly suggestive of there being a huge loss of clonotypes during contraction to their nadirs, followed by compensatory oligoclonal expansions driving a loss of diversity. Conversely those treated with photons underwent far fewer significant remodeling changes, losing fewer T cells and TCRs, retaining more rearrangements over the course of the follow up. It even appears that while proton-treated patient T cell compartments remain stable, the more dramatic reshaping observed in photon treatment appears to have brought those patients’ repertoires in to line with proton group. Arguably, these patients’ T cell compartments might be considered to have been prematurely ‘aged’ (i.e. decreasing naïve T cell frequencies, increasing the proportion of differentiated T cells, decreasing diversity). The extent of the loss of TCR diversity is particularly noteworthy, reaching nadirs that are similar in magnitude to the CD4-depleted and CD8-expanded repertoires we previously observed in untreated chronically-infected HIV patients^38^.

Through correlation of the changes in different T cell populations and the corresponding changes in TCR diversity, it seems that a large increase in the frequency of terminally differentiated CD8+ Temra cells is helping drive this loss of diversity in photon-treated patients. While TCR clustering with known specificity receptors identified potential post-nadir expansion of cytomegalovirus-reactive clones in two patients, potentially due to viral reactivation due to the loss of viral control during the radiation-induced lymphopenia^39^, these particular donors did not see large post-nadir diversity shifts and thus the antigens responsible remain unknown. Regardless, such large oligoclonal expansions reduce the evenness of a repertoire. Altered repertoire diversity has demonstrated potential as a diagnostic tool^40^, however radically decreased diversity may play a more clinically relevant role. Having fewer distinct clones in circulation theoretically makes an immune system less able to respond to as broad an array of antigens, as the likelihood of a presented antigen being recognized by a suitable receptor decreases. Indeed a recent study employing stereotactic body radiotherapy (SBRT) to treat non-small-cell lung cancer reported that patients who developed metastases after treatment had significantly less diverse, more oligoclonal TCR repertoires^41^. In the context of ICB, it’s possible that some T cells that might otherwise be able to respond to presented neoantigens instead die from irradiation before they were activated to kill tumor cells.

We also observed another photon-treatment specific alteration that could be deleterious to ICB: Treg cell frequency rose markedly, across both two post-baseline samples. This is line with previous findings: there are mouse models in which Tregs expand following radiotherapy^42,43^, and clinical data demonstrating the same in patients^44,45^, potentially as a function of both relative Treg radioresistance and increased production. These additional inhibitory cells could provide an additional hurdle for ICB to overcome in order to successfully release anti-cancer immune responses.

It is also possible that the different forms of radiation therapy differentially alter other immune parameters (known to be affected by photon-treatment) which were not studied here. This could include: the production of different cytokines and chemokines^46^, alterations to the immunopeptidome and amount of MHC expressed^47,48^, and DNA damage leading to both local inflammation and *de novo* neoantigen production^49,50^.

Many of these changes either potentially could or are (in some cases) known to synergize with ICB, driving the abundance of combination trials currently ongoing, but it’s possible that these benefits are being blunted by merit of destruction, exhaustion, or suppression of potentially responding clones. Moreover, radiation-induced lymphopenia itself correlates with poorer prognosis and shorter survival times^6^, which is reason enough to try to understand and mitigate its risks. Indeed, in our cohort we saw a correlation between the change in diversity on treatment and the overall survival time from diagnosis, with those patients whose TCR repertoires remaining stably polyclonal in the face of radiotherapy surviving longer. Larger, better-powered cohorts will be required to see if this associations holds true.

As more proton beam centers are constructed, and trials continue to increase the breadth of cancers that might be treatable using protons, the field should ensure it also measures immune parameters as potential correlates of protection. Regardless of radiation type, it is also possible that treatment alterations or additional interventions could be introduced to reduce the impact upon T cells and lymphoid tissues, such as the ‘As Low As Reasonably Achievable’ dosing strategy for lymphocyte-rich tissues as has recently been proposed^51^. Such lymphocyte-sparing radiation might be expected to leave a greater portion of the T cell repertoire in place to respond against cancer antigens once unleashed by immunotherapy.

## Acknowledgements

The authors would like to thank all of the patients and their families for contributing samples to this study.

## Data and Code Availability

Primary TCR sequencing data available from the immuneACCESS resource from https://clients.adaptivebiotech.com/pub/heather-2020. Secondary AIRR-seq community formatted data are available on Zenodo via the DOI <u>10.5281/zenodo.11480289</u>. All scripts and metadata used to generate the plots in this study are available on GitHub from the URL https://github.com/JamieHeather/radiation-induced-lymphopenia-paper-analysis.

## Author Contributions

JMH: Data analysis, drafted manuscript, experimental design

DWK: Data collection, clinical care, experimental design

SMS: Wet lab/flow cytometry

EVS & MGF: Clinical coordination, data collection

TH: Manage cohort, clinical care

RC, NH, TH, MC: Experimental design, project oversight, sourced funding

All authors: Contributed to and critically appraised manuscript

## Conflicts of interest

No potential conflicts of interest were disclosed.

## Notes

https://clients.adaptivebiotech.com/pub/heather-2020

https://doi.org/10.5281/zenodo.11480289

https://github.com/JamieHeather/radiation-induced-lymphopenia-paper-analysis

### Competing Interest Statement

The authors have declared no competing interest.

