## Supplementary Figures for "The differential immunological impact of photon vs proton radiation therapy in high grade lymphopenia"

S Fig 1: Cancer type distribution of radiation-naïve chemoRT patients

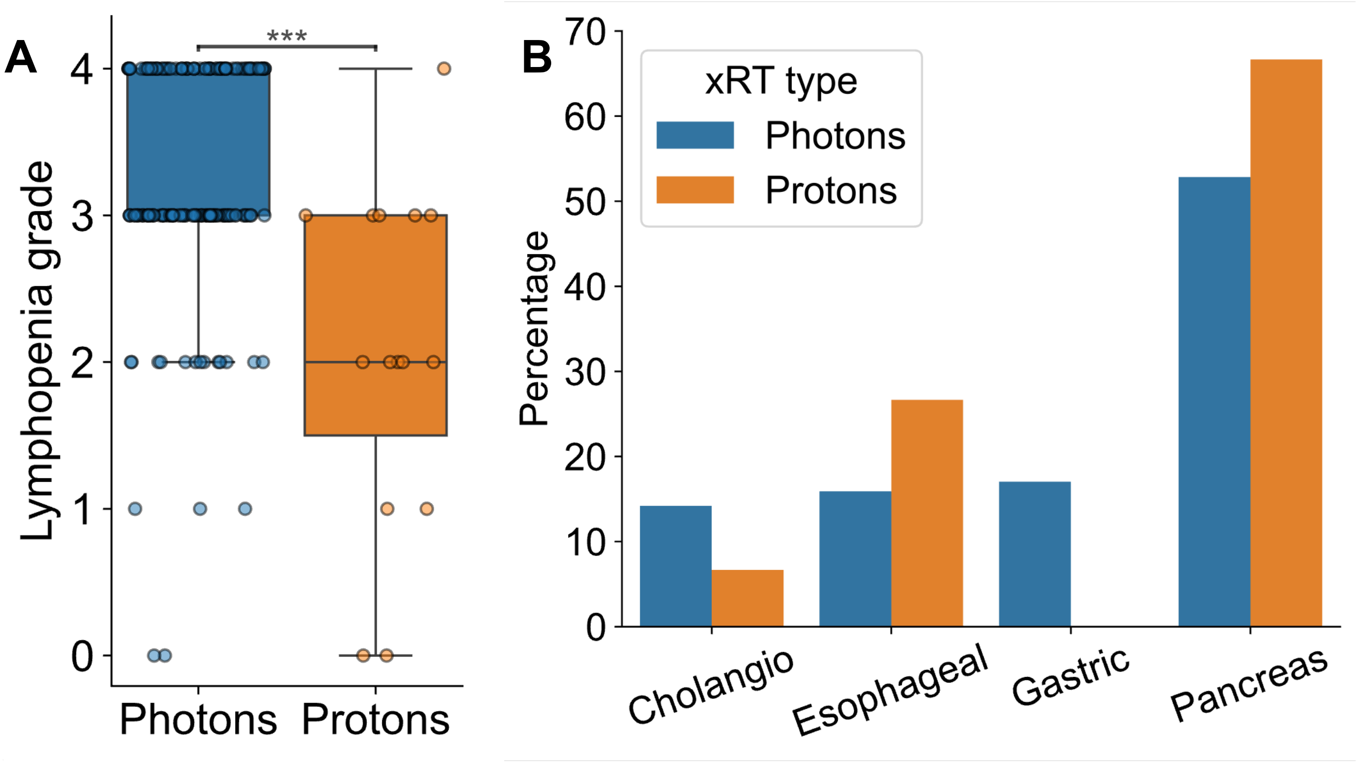


**A**: Lymphopenia grades of of our wider radiation-naïve patient cohort at their respective nadirs in our wider cohort (n = 190 patients, 175 who received photons and 15 who received protons). Boxes show the interquartile range and median, overlain with the individual markers. ***P < 0.001, Mann Whitney U test. .

**B**: Distribution of photon- and proton-treated patients in the previously radiation-naïve subset of the cohort, whose chemoRT-induced lymphopenias are detailed in A and in Figure 1.

### S Fig 2: Cohort composition

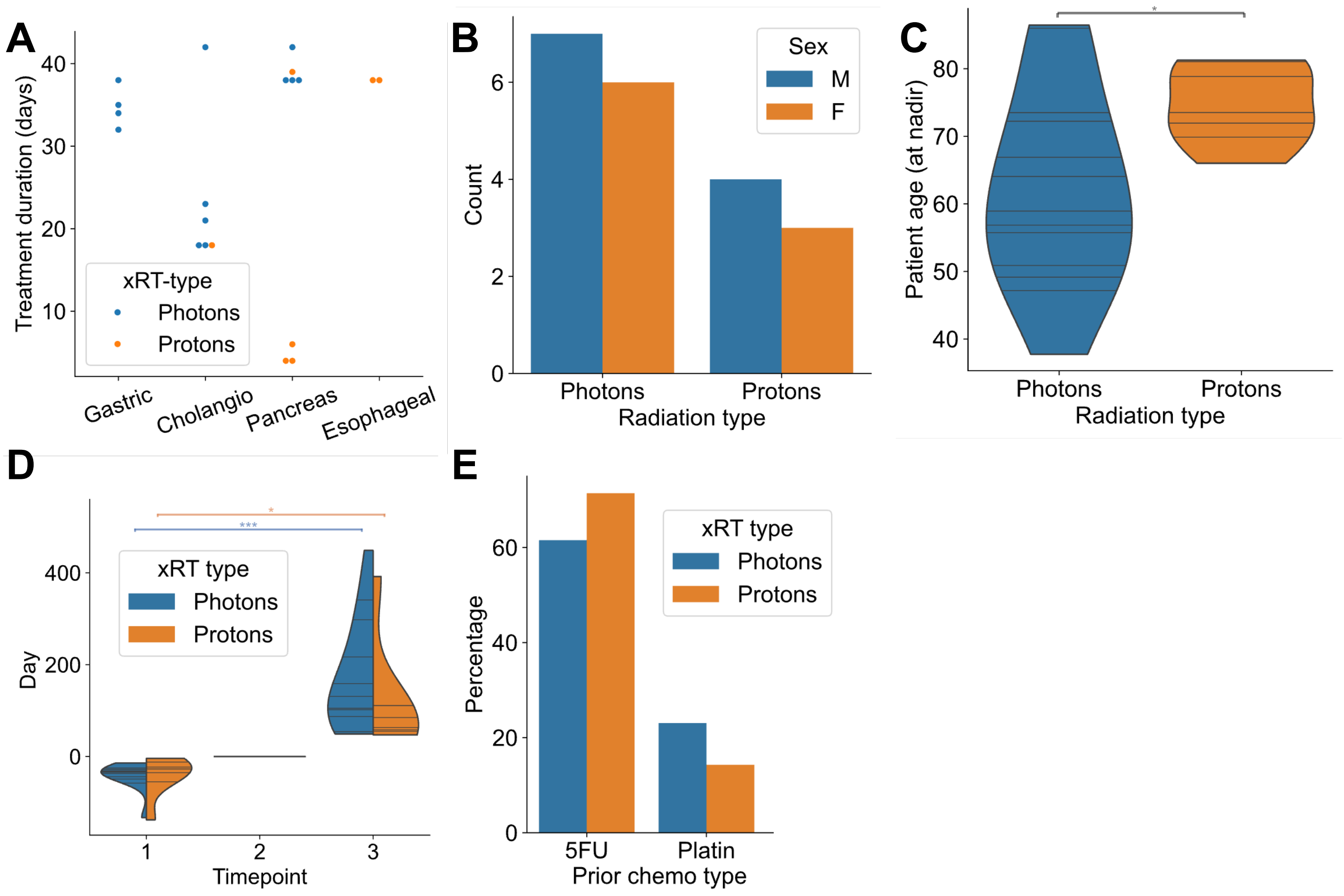


**A**: Duration of xRT (from start to end date) of each patient, colored by treatment type.

**B**: Number of patients of each sex across the two xRT groups.

**C**: Patient ages across the two xRT groups (at the time of their nadir sample time point). The two distributions are significantly different (Mann Whitney U test, P=0.029). Horizontal lines indicate patient values, violin shape indicates kernel density estimations (cut at the terminal observed values).

**D**: Distribution of timepoints in days, made relative to each donor's nadir timepoint. I.e. typically the baseline timepoint (1) was approximately 30 days prior to the nadir (-30), while the recovery sample was typically >= 100 days after (+100).

**E**: Barchart counts of which patients received what prior chemotherapy conditioning regimen. '5FU' indicates patients who received a FOLFOX or FOLFIRINOX based treatment, while 'Platin' indicates those who received a platinum based treatment (including Cisplatin or Carboplatin).

### S Fig 3: Gating strategy for flow panel #1

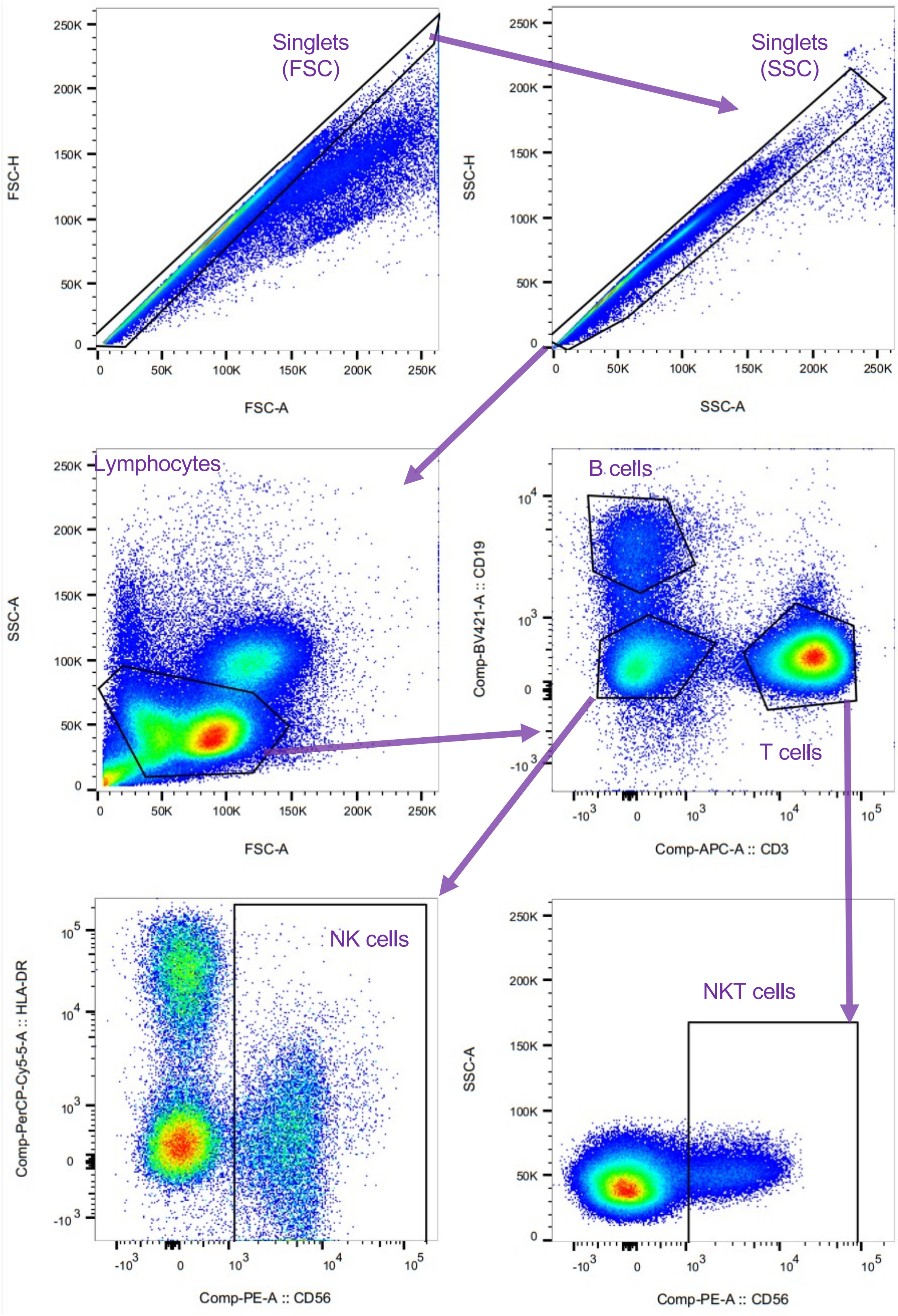


Gating strategy for flow panel 1, after antibodies whose epitopes didn't survive fixation (or those whose utility relied on those which failed) have been removed. Example from normal donor PBMC.

### S Fig 4: Gating strategy for flow panel #2

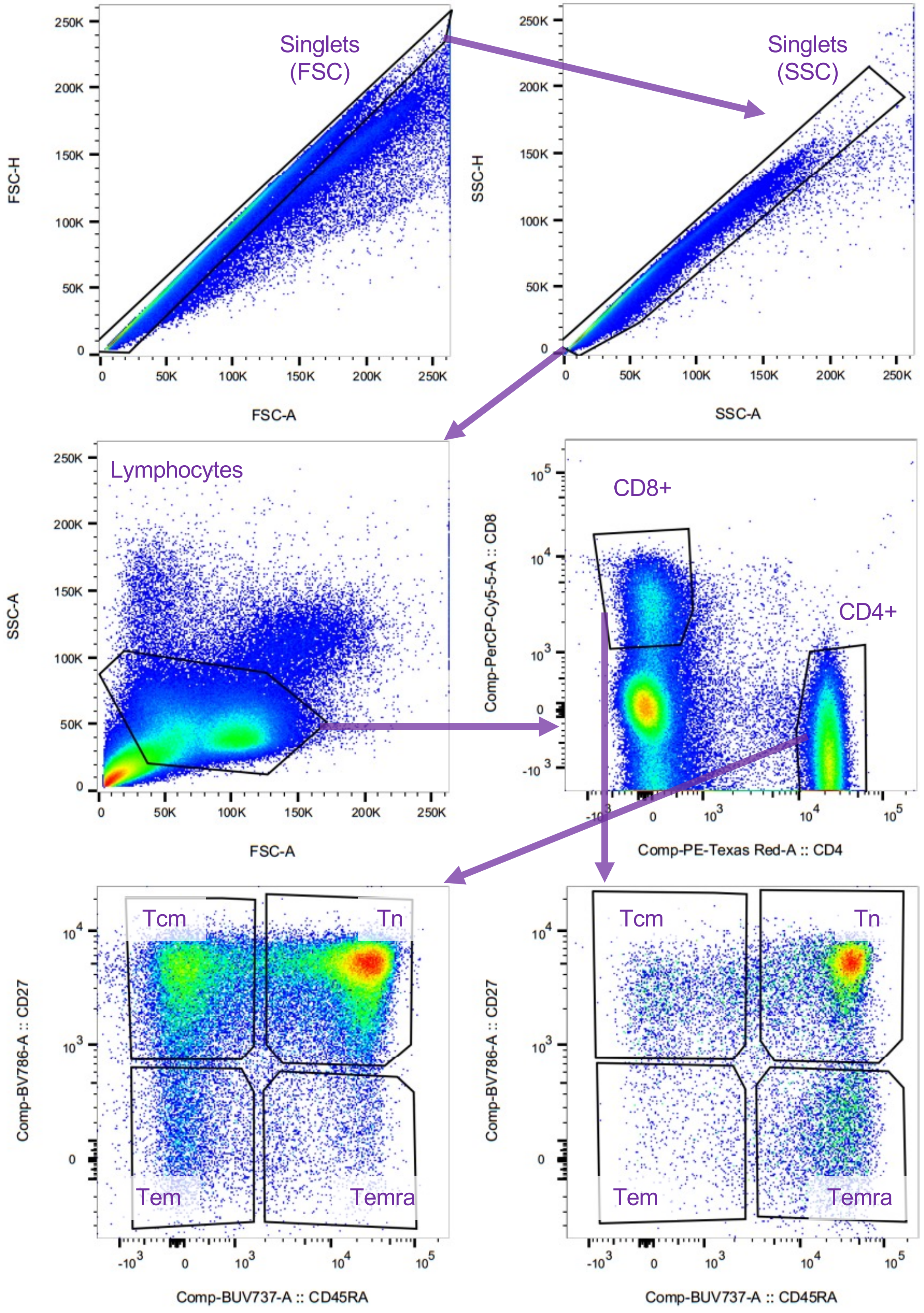


Gating strategy for flow panel 2, after antibodies whose epitopes didn’t survive fixation (or those whose utility relied on those which failed) have been removed. Example from normal donor PBMC.

### S Fig 5: Gating strategy for flow panel #3

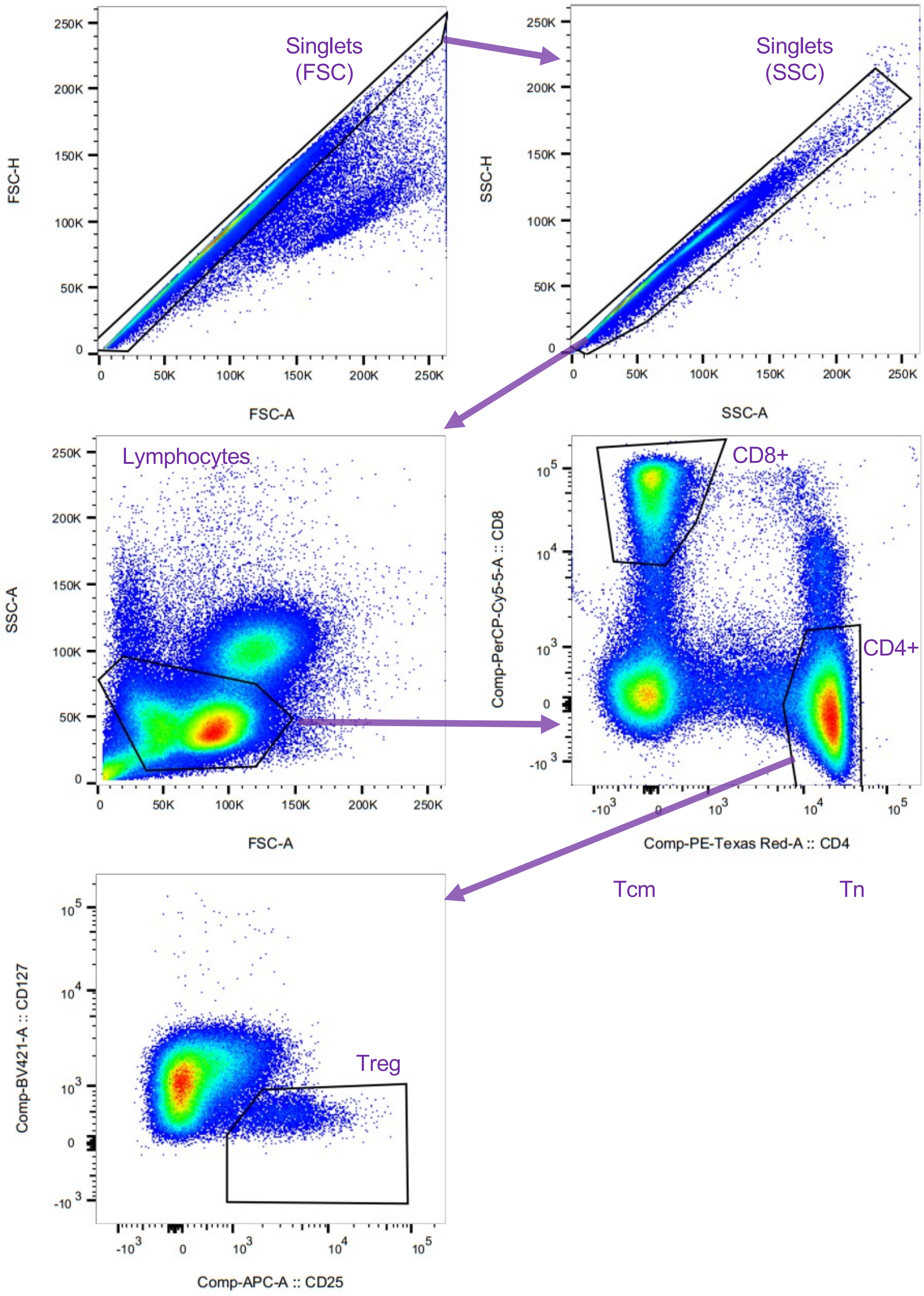


Gating strategy for flow panel 3, after antibodies whose epitopes didn't survive fixation (or those whose utility relied on those which failed) have been removed. Example from normal donor PBMC.

### S Fig 6: Fixed/frozen blood cell flow cytometry sanity checks

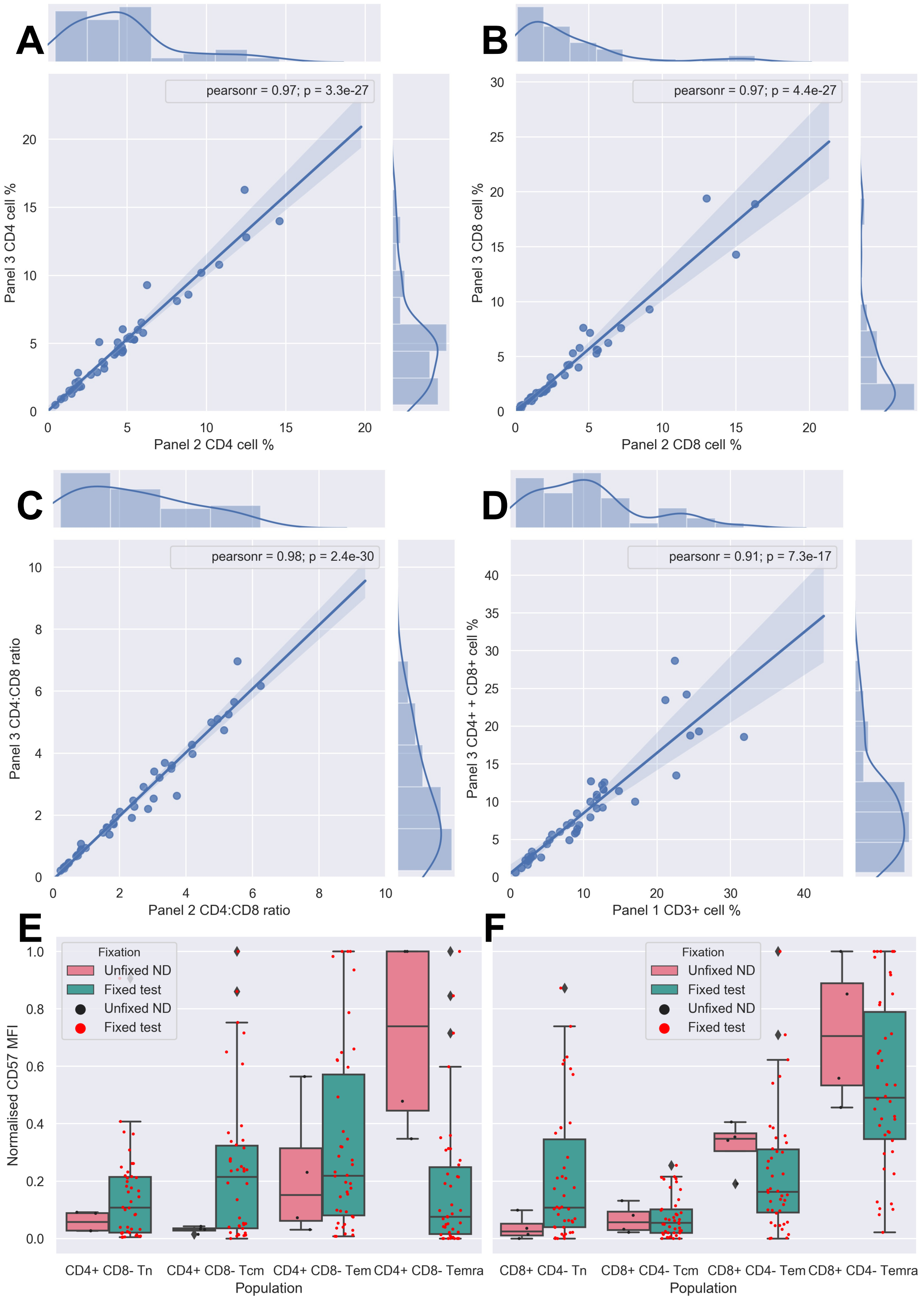


**A & B**: Jointplots showing the relationship between the same antibody stain in different panel 2 (x) and panel 3 (y), for CD4 and CD8 staining respectively. Inset graph shows linear regression between the two (shaded area being 95% confidence intervals), edge histograms show the distribution of the data of each axis individually.

**C**: Jointplot showing the relationship between CD4:CD8 ratio of panel 2 and panel 3.

**D**: Jointplot showing the relationship between panel 1 CD3 % and the sum of panel 3 CD4 and CD8 %s.

**E & F**: Box and whisker plots overlain with stripplots showing the distribution of normalized MFI for CD57 expression for the different T cell subpopulations determined by CD45RA/CD27 expression (see Supplementary Figure 4) for CD4+ (**E**) and CD8+ (**F**) cells. Central lines indicate medians, boxes show inter-quartile range (IQR), whiskers show IQR * 1.5. Fixed radiotherapy patient samples in green, three unfixed normal donor leukopak samples shown in pink as a reference.

### S Fig 7: B cell frequencies in xRT patients across treatment

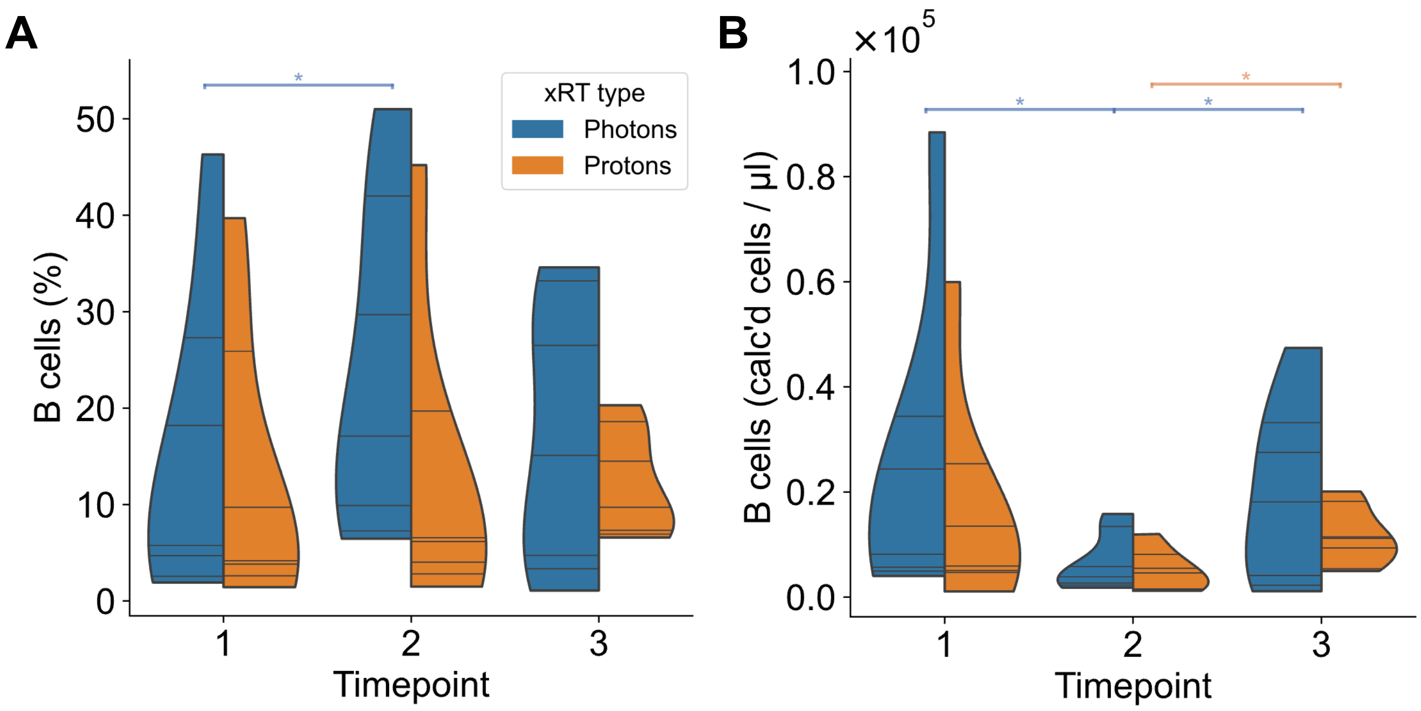


**A**: Violin plots of the percentage of CD19+ B cells in blood samples of of photon (blue) and proton (orange) treated cancer patients at each of the three time points. Horizontal lines indicate patient values, violin shape indicates kernel density estimations (cut at the terminal observed values). Black significance lines indicate intra-time point unpaired non-parametric tests (Mann Whitney U), while blue and orange lines indicate inter-time point paired non-parametric tests (Wilcoxon ranked-sum). *P < 0.05, **P < 0.01.

**B**: As in A, but showing calculated absolute cell numbers, using the percentage values combined with the corresponding calculated absolute lymphocyte counts.

### S Fig 8: T cell differentiation status changes over radiation therapy

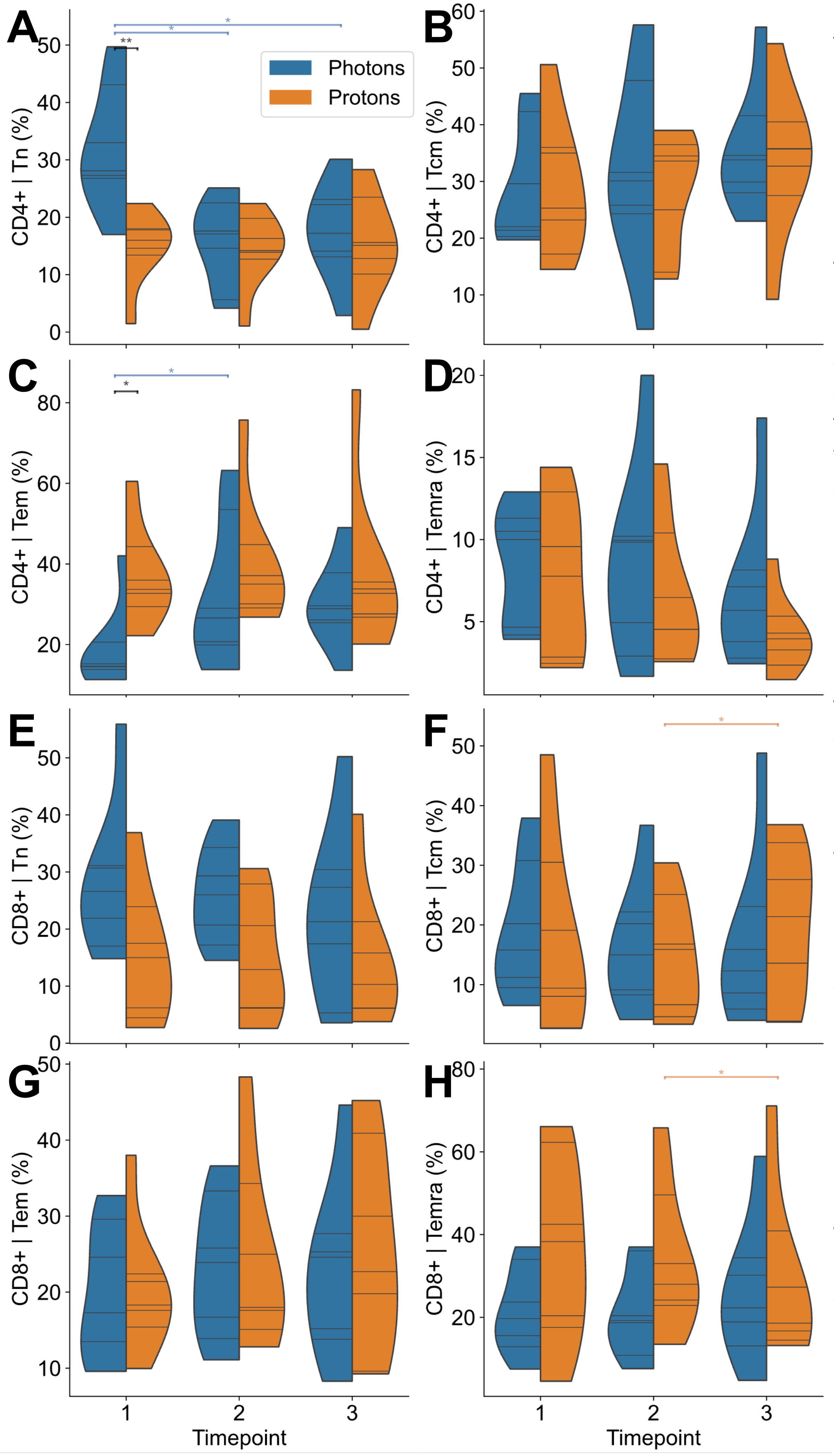


Violin plots of the percentage of different T cell subpopulations in blood samples of photon (blue) and proton (orange) treated cancer patients at each of the three time points. Horizontal lines indicate patient values, violin shape indicates kernel density estimations (cut at the terminal observed values). Black significance lines indicate intra-time point unpaired non-parametric tests (Mann Whitney U), while blue and orange lines indicate inter-time point paired non-parametric tests (Wilcoxon ranked-sum). *$P < 0.05$, **$P < 0.01$. **A**: Naïve CD4+ T cells (CD27+ CD45RA+). **B**: CD4+ central memory T cells (CD27+ CD45RA-). **C**: CD4+ effector memory T cells (CD27- CD45RA-). **D**: Terminally differentiated CD4+ effector memory T cells (CD27- CD45RA+). **E**: Naïve CD8+ T cells (CD27+ CD45RA+). **F**: CD8+ central memory T cells (CD27+ CD45RA-). **G**: CD8+ effector memory T cells (CD27- CD45RA-). **H**: Terminally differentiated CD8+ effector memory T cells (CD27- CD45RA+).

### S Fig 9: Subsampled TCR repertoire diversity metrics

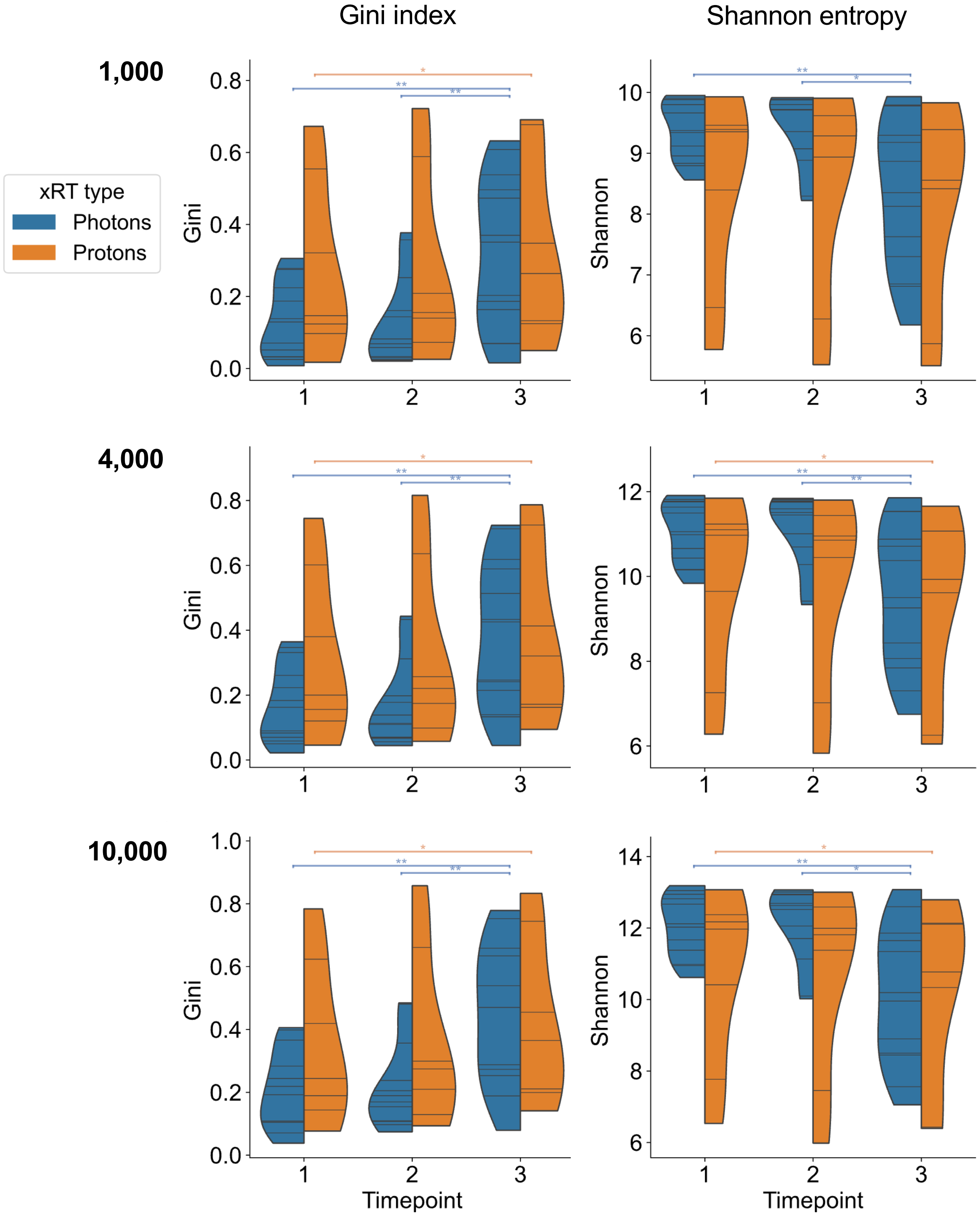


Randomly size-matched sub-sampled data for Gini Index and Shannon Entropy, as shown in Figure 4. A set number of TCRs were randomly drawn from each repertoire (1000, 4000, or 10000) 100 times, diversity metrics calculated at each iteration, then mean values calculated and plotted. Violin area shapes indicate kernel density estimations (cut at the terminal observed values). Blue and orange lines indicate inter-time point paired non-parametric tests (Wilcoxon ranked-sum). *$P$ < 0.05, **$P$ < 0.01.

### S Fig 10: Changes in TCR repertoire diversity metrics between timepoints

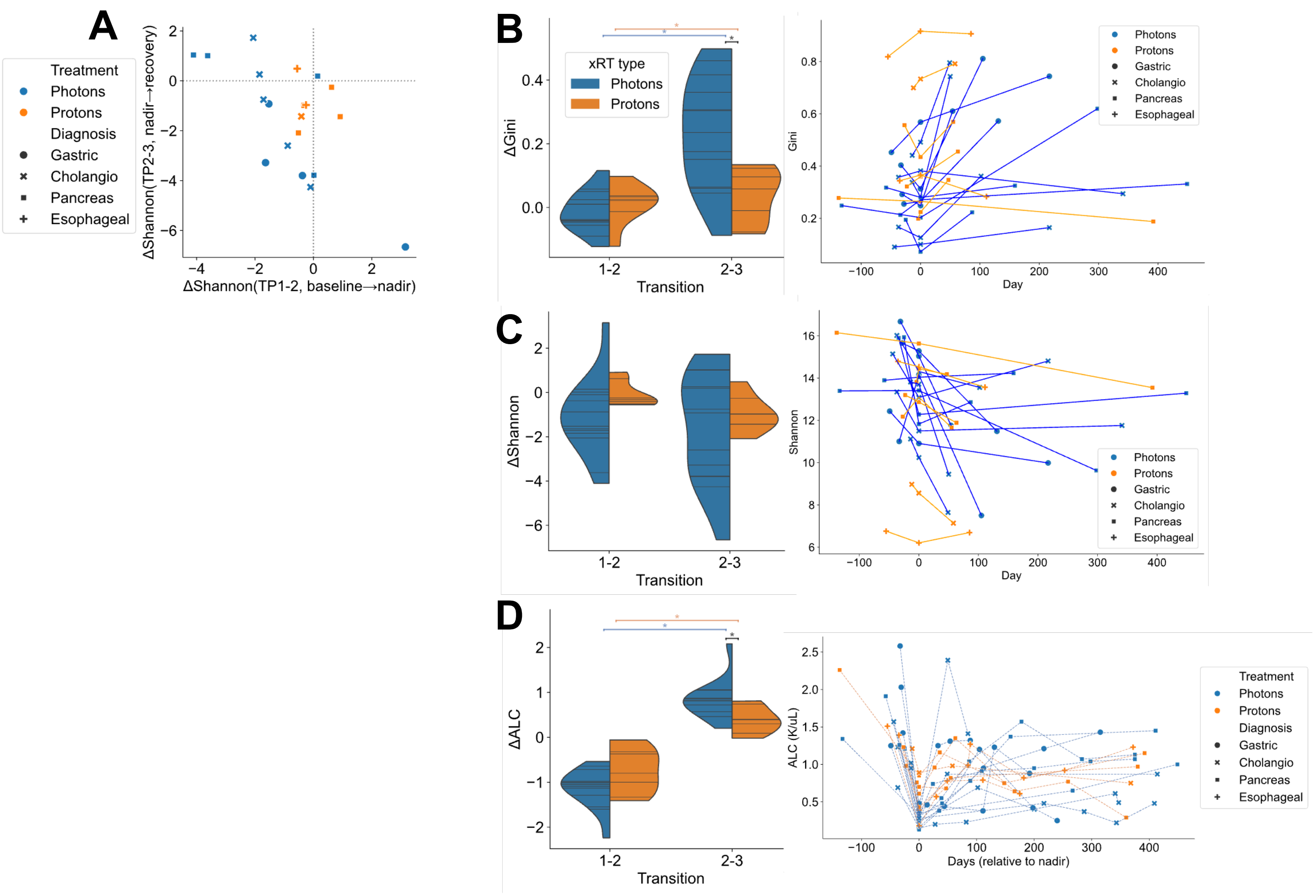


**A**: Scatterplot of the change in Shannon Entropy of each patient from between timepoints 1 and 2 (x axis) and 2 and 3 (y axis). Samples are colored by treatment type, and markers are assigned by diagnosis.

**B**: Violinplot of the distribution of changes in Gini Index between timepoints 1 and 2, and 2 and 3 (left). Violin area shapes indicate kernel density estimations (cut at the terminal observed values). Black significance lines indicate intra-time point unpaired non-parametric tests (Mann Whitney U), while blue and orange lines indicate inter-transition paired non-parametric tests (Wilcoxon ranked-sum). $^*P < 0.05$, $^{**}P < 0.01$. Right line chart shows the actual values plotted by day (relative to nadir).

**C**: As in **B**, but for Shannon Entropy.

**D**: As in **B**, but for ALC. Note that line chart on right goes to later timepoints, possible due to additional clinical samples from which ALC was measured but no material was available for other assays.

### S Fig 11: Inter-donor sample overlap of subsampled TCR repertoires

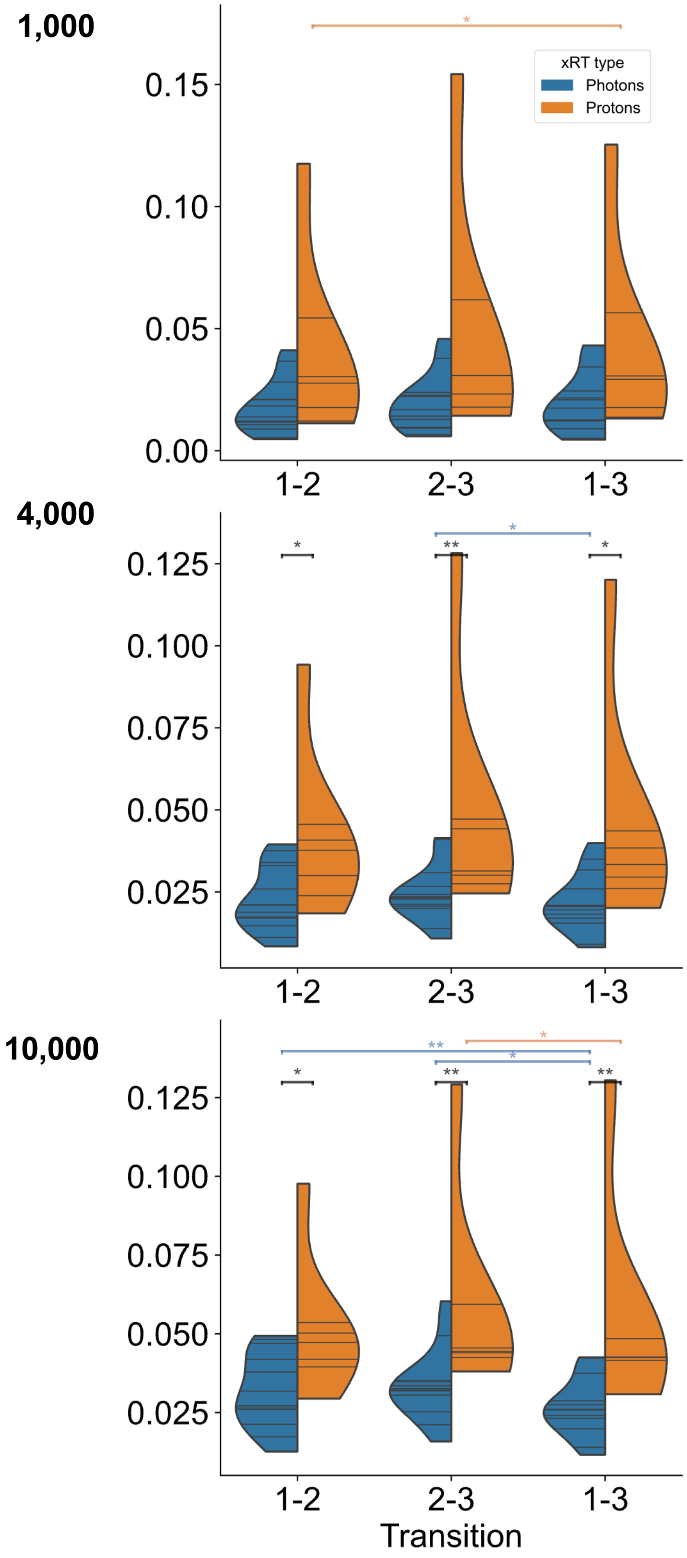


Randomly size-matched sub-sampled TCR data was used for Jaccard Index calculation, as shown in Figure 4F. A set number of TCRs were randomly drawn from each repertoire (1000, 4000, or 10000) 100 times, Jaccard indices were calculated for each transition, then finally all 100 iterations were averaged and plotted. Violin area shapes indicate kernel density estimations (cut at the terminal observed values). Blue and orange lines indicate inter-transition paired non-parametric tests (Wilcoxon ranked-sum). **P* < 0.05, ***P* < 0.01.

S Fig 12 : Baseline cell populations correlate with repertoire diversity

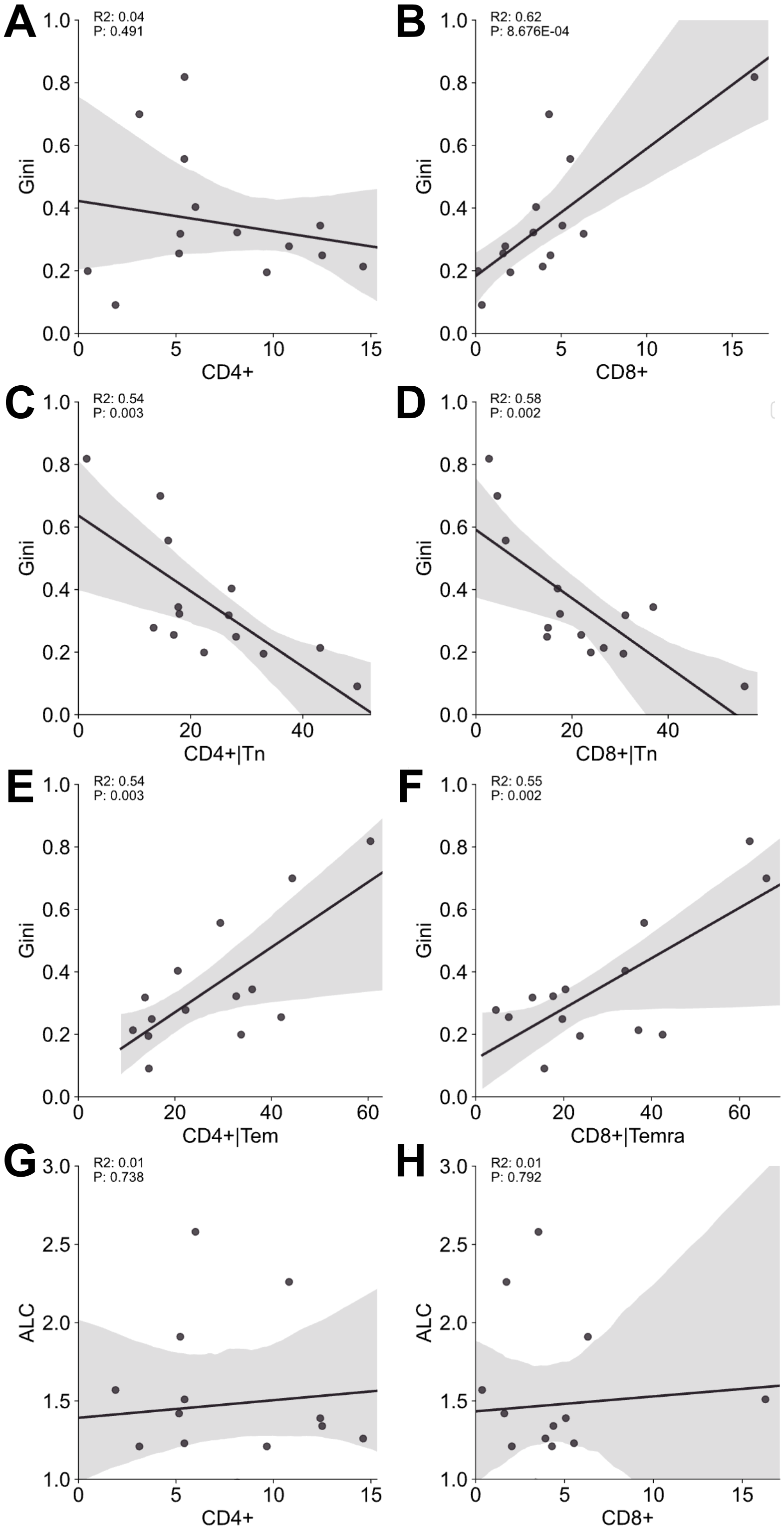


**A-F**: Linear regressions between various flow cytometric population frequencies (%) for all patient samples at baseline (prior to treatment) and the Gini index of the corresponding subsampled TCRb repertoires. 4000 TCRs were subsampled 100 times and Gini indexes averaged. X axes show: overall CD4+ T cell % (**A**); overall CD8+ T cell % (**B**); CD4+ naïve T cell % (**C**); CD8+ naïve T cell % (**D**); CD4+ effector memory T cell % (**E**); and CD8+ Temra cell % (**F**).

**G-H**: Similar linear regression of overall CD4+ and CD8+ T cell frequencies respectively with ALC, showing a lack of correlation.

### S Fig 13 : CD8+ naïve and Temra T cell frequencies versus Shannon entropy.

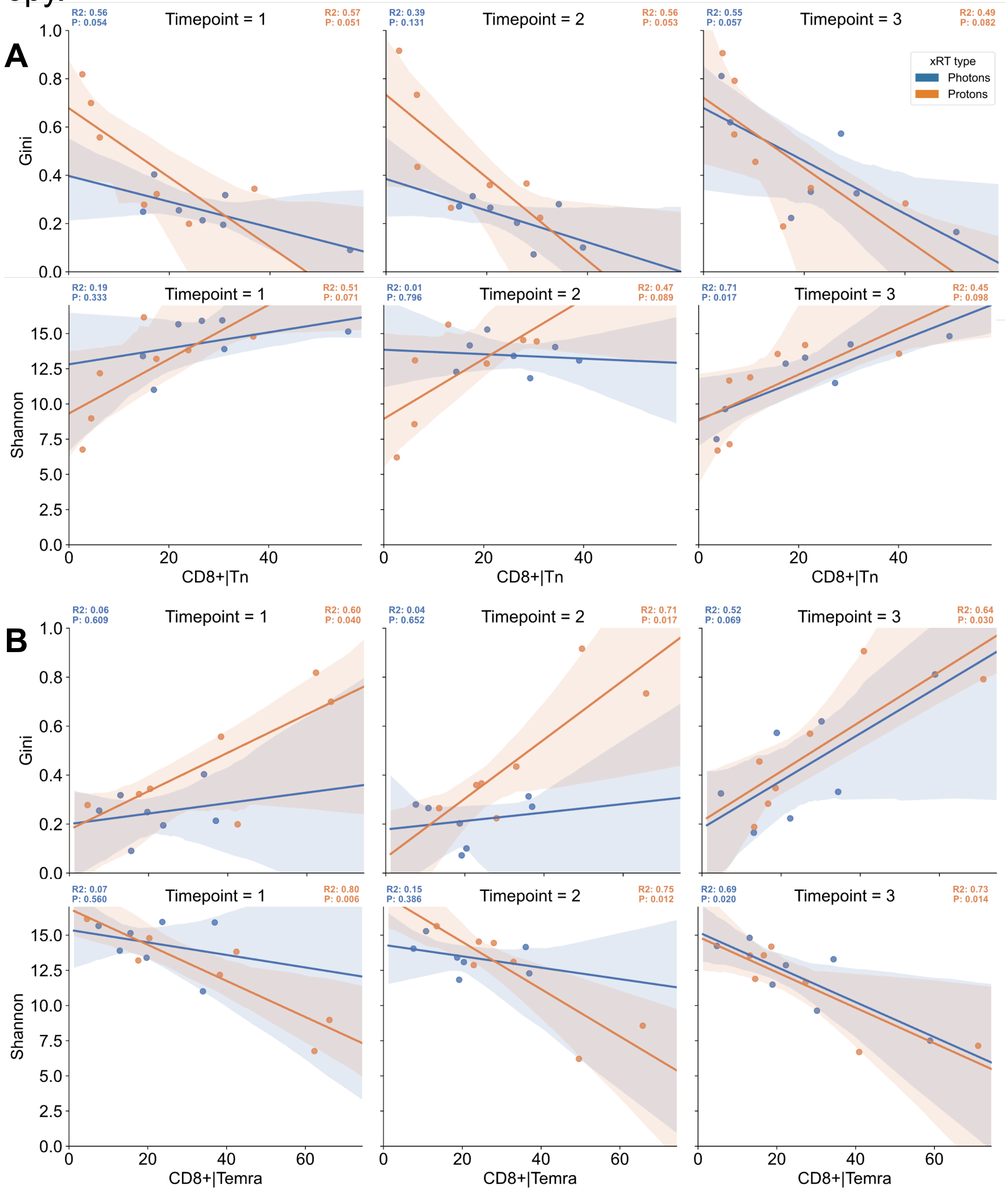


**A & B**: Linear regression of the photon (blue) and proton (orange) treated patient samples, showing percentage frequency of naïve CD8+ T cells (**A**) or CD8+ Temra cells (**B**) on the x axis and Gini index (upper) and Shannon entropy (lower) of size-matched TCR repertoires (average of sampling 4000 TCRs 100 times) on the y, for each time point. Shaded areas indicate 95% confidence intervals. Color-matched $R^2$ and P values displayed above show each patient group's regression statistics.

### S Fig 14 : CD4+ T cell populations correlation with diversity metrics

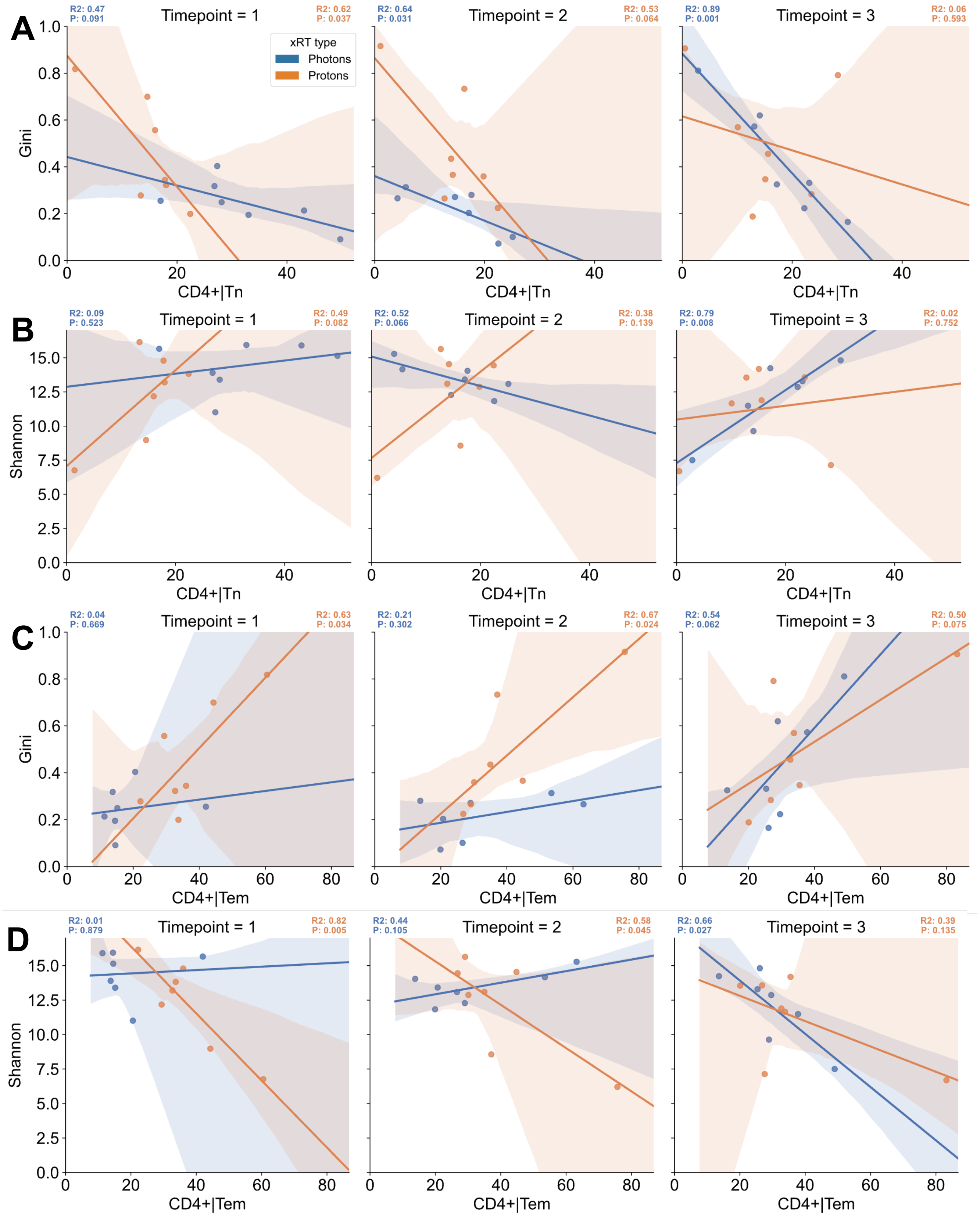


**A & B**: Linear regression of the photon (blue) and proton (orange) treated patient samples, showing percentage frequency of naïve CD4+ T cells on the x axis and either Gini index (**A**) or Shannon entropy (**B**) of size-matched TCR repertoires (average of sampling 4000 TCRs 100 times) on the y, for each time point. Shaded areas indicate 95% confidence intervals. Color-matched $R^2$ and P values displayed above show each patient group's regression statistics.

**C & D**: As with **A** & **B**, save for CD4+ Tem cells on the x axes.

### S Fig 15 : Change in CD8+ Temra frequency correlates with change in Shannon entropy

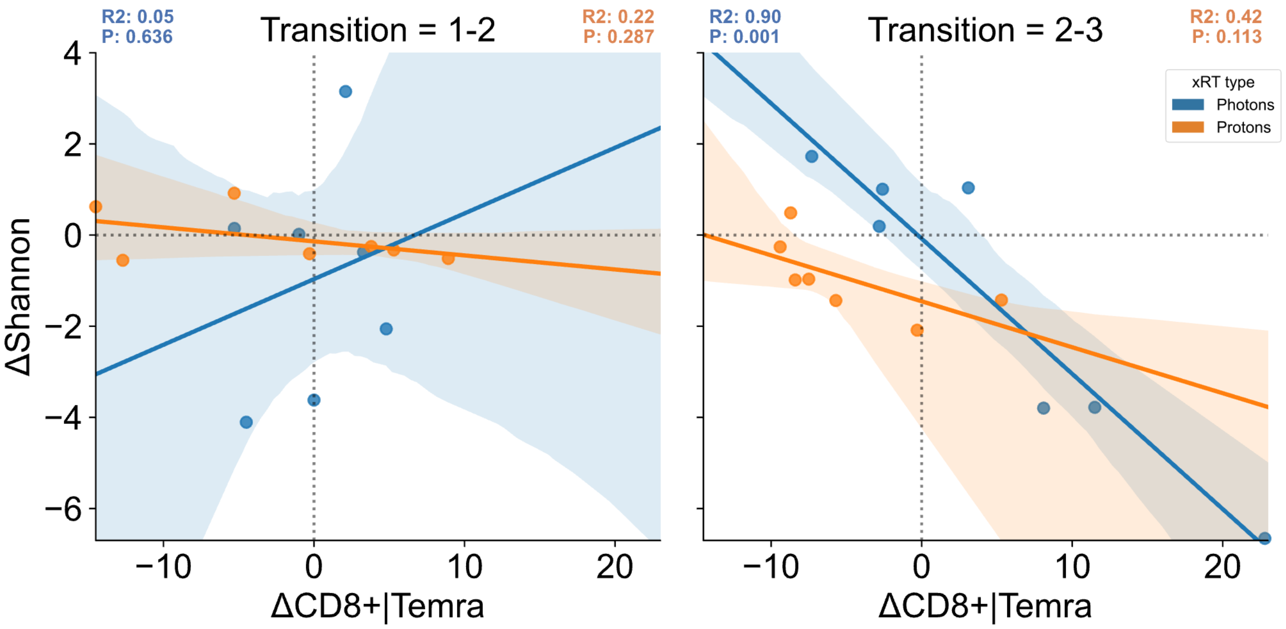


Linear regression showing the change in values between two blood draws; on the left, timepoint 1 to 2 (baseline to nadir), and timepoint 2 to 3 (nadir to recovery) on the right. Changes expressed as a delta, with the first value subtracted from the latter. X axis shows the change in CD8+ Temra cell % frequency for the two transitions, while the y axis shows the change in Shannon entropy for each.

### S Fig 16 : Clonal dynamics for the top 100 rearrangements per time point.

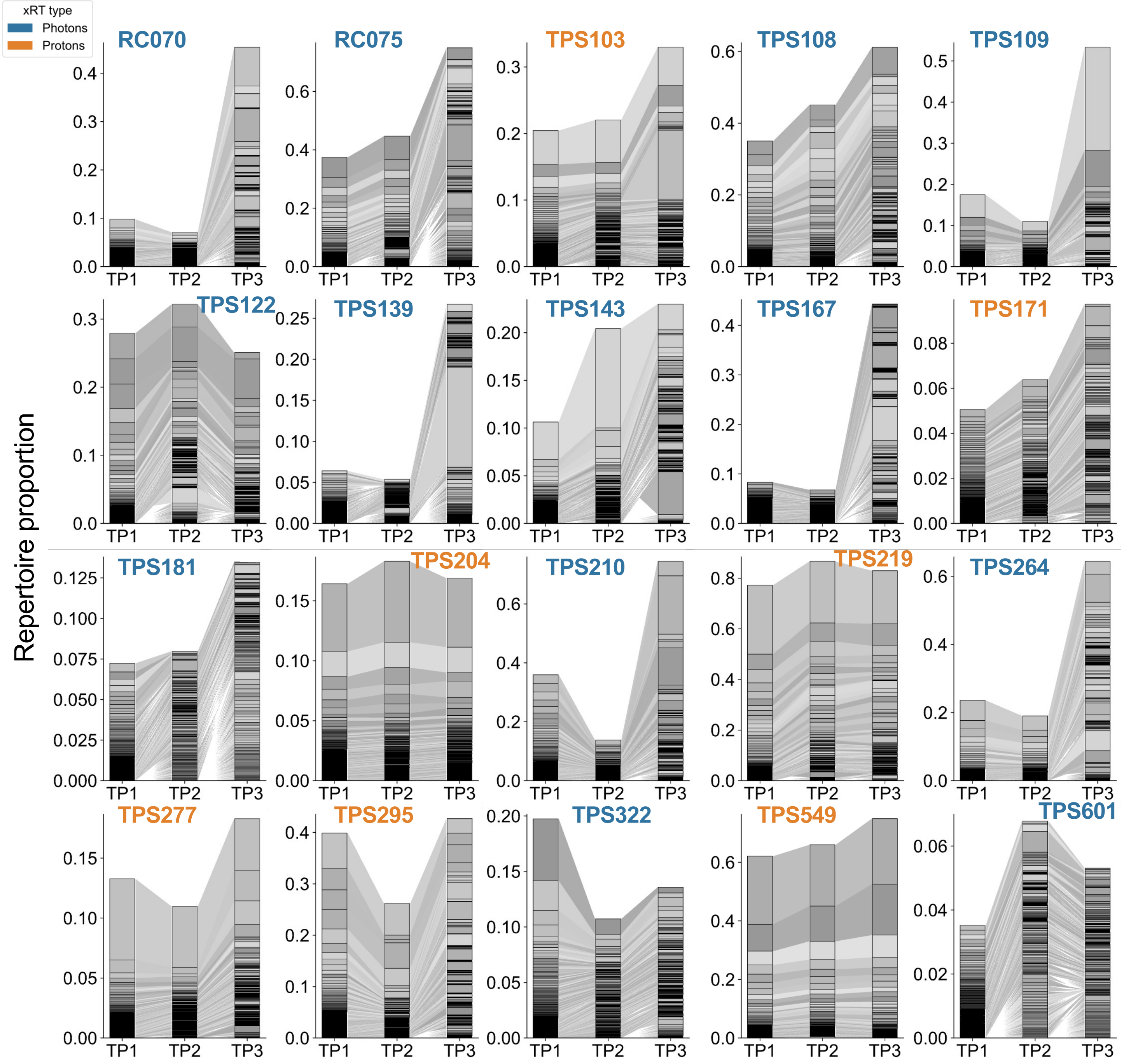


**Above**: Sankey-style flow plots showing the change in frequency of the most abundant TCR rearrangements in every donor, colored by donor . The top 100 rearrangements per donor per time point were pooled, assigned random greyscale colors, and plotted in stacked bar-charts with connecting shaded areas (with absence in a timepoint indicated by shaded areas originating from the halfway point between stacks), sorted by their abundance in TP1.

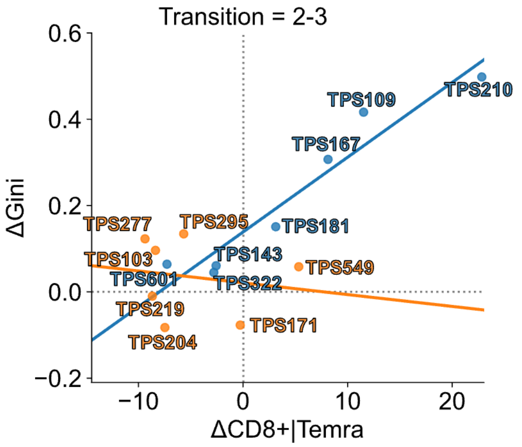


**Right**: alternative plot of Figure 4A right hand panel, showing the patient identifiers used above.

### S Fig 17 : Potential antigenic classifications by TCR clustering.

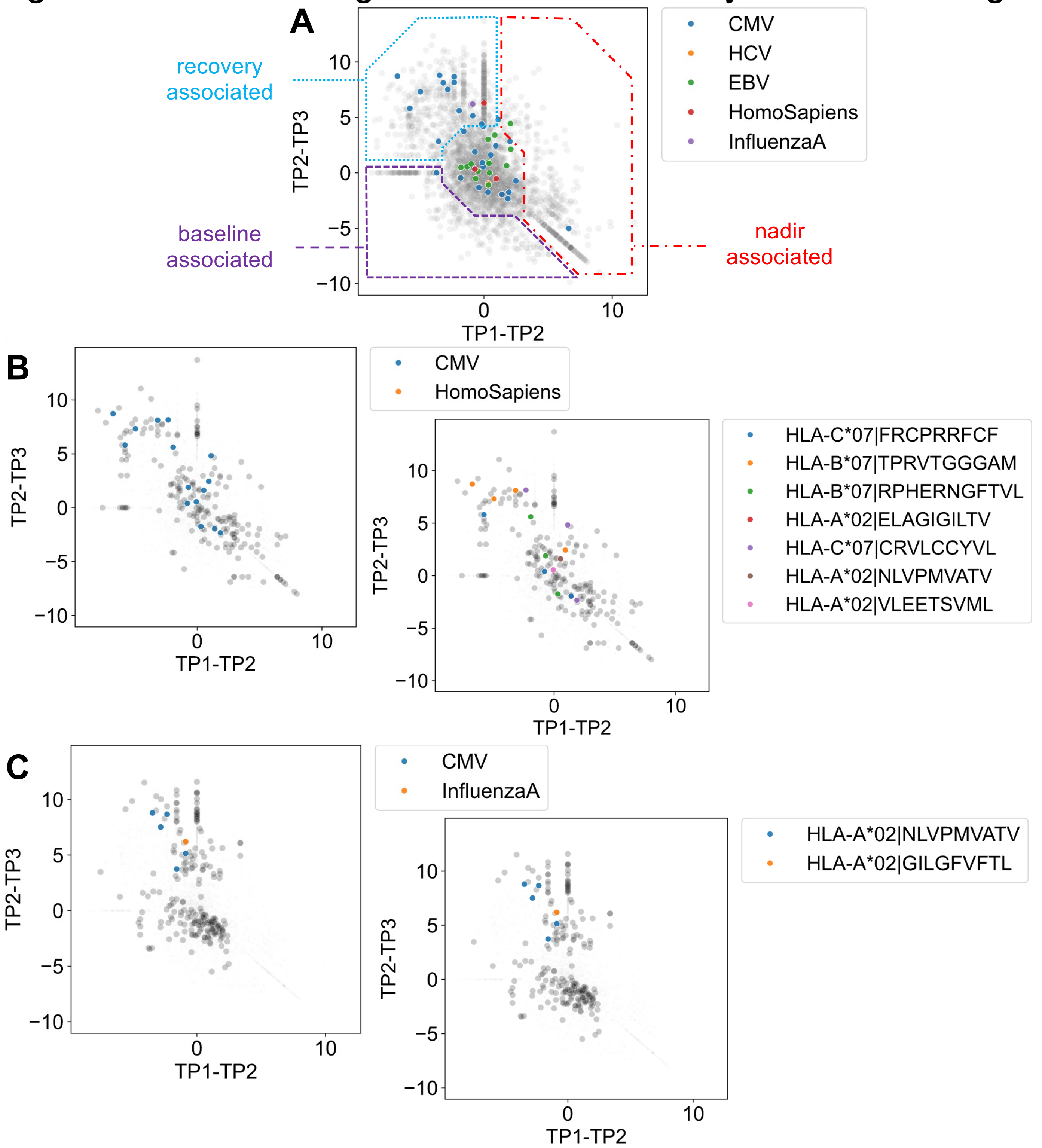


**A**: Potential antigen specificities predicted by TCR clustering for TCRs present in the top 100 most abundant clones of every time point of each donor (see the ‘TCR antigen clustering’ section of the Methods). In order to visualize the change in frequency of each TCR across the three points, we’ve plotted the clonal ‘trajectories’, by plotting the change in frequency of each clone between baseline and nadir (TP1-TP2) on the x axis, and between nadir and recovery (TP2-TP3) on the y axis. TCRs not present in a given sample were arbitrarily assigned a frequency half that of the lowest frequency for that donor. All donors plotted here, with rearrangements lacking a predicted specificity in greyscale and species of predicted epitope colored as per legend. Colored borders added to illustrate: baseline-associated TCRs (purple/dashed); recovery associated (blue/dotted); nadir-associated (red/dot-dashed).

**B** and **C**: As in **A**, but only showing the TCRs belonging to donors TPS143 and RC070 respectively, with specific epitopes inset right.

### S Fig 18 : post-nadir diversity change survival analysis

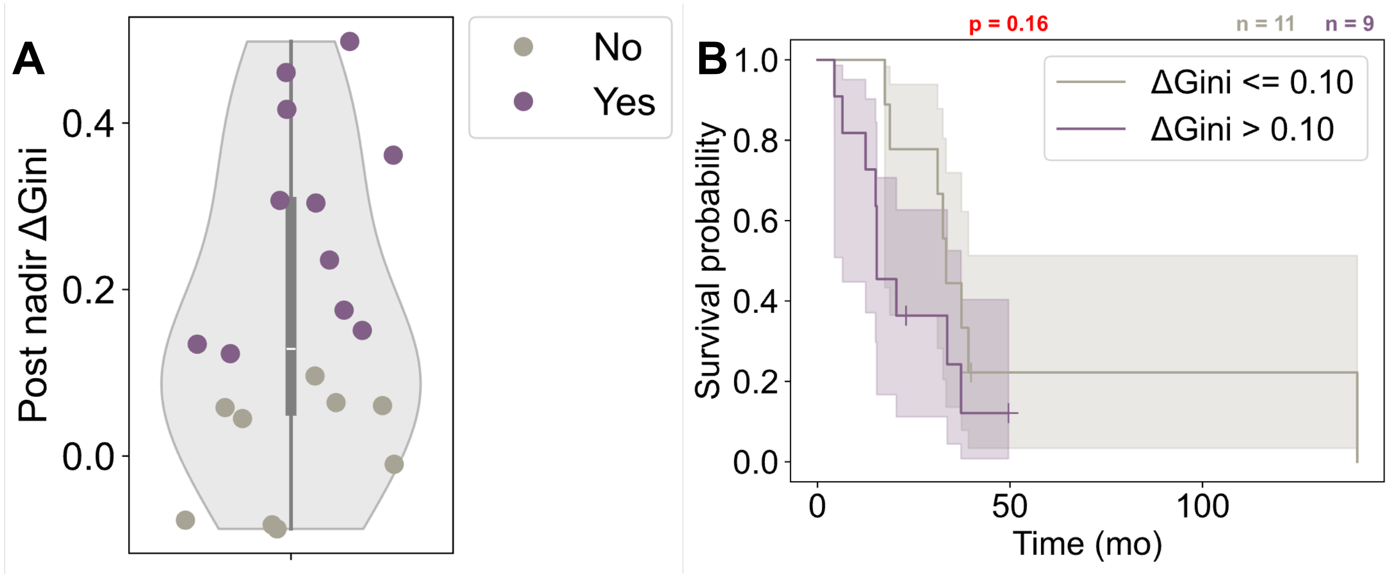


**A**: Violinplot showing he cohort was split in two based on their post-nadir change in Gini index, with those that did undergo a noted increase in oligoclonality (Gini index increase >= 0.1, purple markers), or did not (Gini index change < 0.1, gray markers). **B**: Kaplan-Meier plot of the survival data of the cohort separated out by post-nadir ΔGini change, showing a non-significant difference in survival between the two groups via univariate Cox proportional hazard model analysis (p = 0.16).
